# Mapping eQTLs With RNA-Seq Reveals Novel SLE Susceptibility Genes, Non-Coding RNAs, and Alternative-Splicing Events That Are Concealed Using Microarrays

**DOI:** 10.1101/076026

**Authors:** Christopher A. Odhams, Andrea Cortini, Lingyan Chen, Amy L. Roberts, Ana Vinuela, Alfonso Buil, Kerrin S. Small, Emmanouil T. Dermitzakis, David L. Morris, Timothy J. Vyse, Deborah S. Cunninghame Graham

**Author notes:** Correspondence: Dr Deborah S. Cunninghame Graham.

## Abstract

Studies attempting to functionally interpret complex-disease susceptibility loci by GWAS and eQTL integration have predominantly employed microarrays to quantify gene-expression. RNA-Seq has the potential to discover a more comprehensive set of eQTLs and illuminate the underlying molecular consequence. We examine the functional outcome of 39 variants associated with Systemic Lupus Erythematosus (SLE) through integration of GWAS and eQTL data from the TwinsUK microarray and RNA-Seq cohort in lymphoblastoid cell lines. We use conditional analysis and a Bayesian colocalisation method to provide evidence of a shared causal-variant, then compare the ability of each quantification type to detect disease relevant eQTLs and eGenes. We discovered a greater frequency of candidate-causal eQTLs using RNA-Seq, and identified novel SLE susceptibility genes that were concealed using microarrays (e.g. *NADSYN1*, *SKP1*, and *TCF7*). Many of these eQTLs were found to influence the expression of several genes, suggesting risk haplotypes may harbour multiple functional effects. We pinpointed eQTLs modulating expression of four non-coding RNAs; three of which were replicated in whole-blood. Novel SLE associated splicing events were identified in the T-reg restricted transcription factor, *IKZF2*, the autophagy-related gene *WDFY4*, and the redox coenzyme *NADSYN1*, through asQTL mapping using the Geuvadis cohort. We have significantly increased our understanding of the genetic control of gene-expression in SLE by maximising the leverage of RNA-Seq and performing integrative GWAS-eQTL analysis against gene, exon, and splice-junction quantifications. In doing so, we have identified novel SLE candidate genes and specific molecular mechanisms that will serve as the basis for targeted follow-up studies.

## Introduction

Genome-Wide Association Studies (GWAS) have successfully identified a large number of genetic loci that contribute to complex-disease susceptibility in humans (1). Evidence suggests these variants are enriched within regulatory elements of the genome and their effects play a central role in modulation of intermediate quantitative phenotypes such gene expression (1–6). Many expression quantitative trait loci (eQTL) mapping studies have since been conducted across a wide-range of ethnicities (7, 8), cell-types (9–16), disease states (17–22) and in response to various environmental stimuli (23, 24) - with each contributing to our understanding of the architecture of human regulatory variation in complex-disease.

In spite of diverse study designs, a significant constraint on the majority of such investigations is the use of 3´-targeted microarrays to profile gene expression. The effects of splicing are less likely to be detected through quantification of pre-defined probes that target common exons of a gene (25) and may explain why only a limited number of susceptibility loci localize to causal eQTL signals (26, 27). Technical limitations of microarrays and noise from the small probe design of exon-arrays, further hinder the accuracy of expression measurements (25, 28–30). RNA-Seq based eQTL mapping studies are beginning to emerge (31, 32) and, although large-scale analysis pipelines are still being streamlined, such types of investigations will greatly increase the likelihood of capturing disease associated eQTLs as quantification of overall gene and independent exon expression, and relative transcript abundance (including novel isoforms and non-coding RNAs) is possible (33–39).

Integrative studies using RNA-Seq to functionally annotate complex-disease susceptibility loci however have been limited (35, 40–44). Moreover, numerous investigations have aimed to explain the functional relevance of susceptibility loci by interrogation of GWAS SNPs themselves in eQTL datasets and simply testing for association with gene expression (45– 47). Such inferential observations should be treated with caution as they may possibly be the result of coincidental overlap between disease association and eQTL signal due to local LD and general ubiquity of regulatory variants (48). This has become particularly important as statistical power in eQTL cohorts grow and availability of summary-level data accession through eQTL data-browsers increases (49–51).

In this investigation, we integrate eQTL data derived from both microarray and RNA-Seq experiments with our GWAS results in Systemic Lupus Erythematosus (SLE [MIM: 152700]); a heritable autoimmune disease with undefined aetiology and over 50 genetically associated loci (52–54). We use summary-level *cis*-eQTL results in lymphoblastoid cell lines (LCLs) taken from the TwinsUK cohort to directly compare the microarray (9) and RNA-Seq (39) results in detecting SLE associated eQTLs along with their accompanying eGenes. We apply a rigorous two-step approach – a combination of conditional (55) and Bayesian colocalisation (56) analysis – to test for a shared causal variant at each locus. We demonstrate the benefits of using RNA-Seq over microarrays in eQTL analysis by identifying not only novel SLE candidate-causal eGenes but also putative molecular mechanisms by which SLE-associated SNPs may act; including differential exon usage, and expression modulation of non-coding RNA. Our investigation was extended to include RNA-Seq expression data in whole blood in order to validate the eQTL signals detected in LCLs and uncover the differences in genetic control of expression between cell-types. Finally, we interrogate the Geuvadis RNA-Seq cohort (35) to identify SLE associated alternative-splicing quantitative trait loci (asQTLs) and highlight the advantages of profiling at various resolutions to detect eQTLs that would otherwise remain concealed. Through functional annotation of SLE associated loci using microarray and RNA-Seq derived expression data, we have supplied comprehensive evidence of the need to use RNA-Seq to detect disease contributing eQTLs and, in doing so, have suggested novel functional mechanisms that serve as a basis for future targeted follow-up studies.

## Results

### Discovery and classification of SLE candidate-causal eQTLs and eGenes

The first part of this study integrated the 39 SLE associated SNPs taken from a recent GWAS in Europeans (**Table 1**) with eQTLs from the TwinsUK gene-expression cohort (n=856) profiled using microarray and RNA-Seq (at both gene-level and exon-level resolutions). To accomplish this, we implemented a two-step pipeline (**Fig. 1**), and subjected the genomic intervals within +/-1Mb of each of the 39 GWAS SNPs to eQTL association analysis against expression quantifications in LCLs (**Table 2**).

**Table 1.** Independent allelic associations at SLE susceptibility loci following meta-analysis with replication study

|  |  |  |  |  |  |  |
| --- | --- | --- | --- | --- | --- | --- |
| **GWAS SNP**<br>**(Tag SNP)** | **GWAS SNP**<br>**Position (hg19)** | **Variant**<br>**Alleles** | **Odds Ratio (CI)** | **P-value** | **GENCODE v10**<br>**Annotation** | **Annotated eGenes**<br>**from microarray LCLs** |
| rs2476601 | 1:114377568 | G/A | 1.43 (1.34-1.53) | 1.10E-28 | *PTPN22* Missense |  |
| rs1801274 | 1:161479745 | G/A | 1.16 (1.11-1.21) | 1.04E-12 | *FCGR2A* Missense | *FCGR2B* |
| rs10912578 (rs844663) | 1:173251856 | G/A | 1.27 (1.22-1.33) | 4.16E-19 | *LOC100506023* Intronic |  |
| rs10753074 (rs1935325) | 1:173346343 | T/C | 1.21 (1.15-1.26) | 5.82E-12 | *LOC100506023* Intronic |  |
| rs17849501 | 1:183542323 | C/T | 2.10 (1.95-2.26) | 3.45E-88 | *NCF2* Synonymous | *SMG7* |
| rs3024505 | 1:206939904 | G/A | 1.17 (1.11-1.24) | 4.64E-09 | 1kb 3' of *IL10* |  |
| rs9782955 | 1:236039877 | C/T | 1.16 (1.11-1.22) | 1.25E-09 | *LYST* Intronic | *LYST* |
| rs268134 | 2:65608363 | G/A | 1.21 (1.15-1.27) | 1.14E-10 | *SPRED* Intronic | *SPRED2* |
| rs2111485 | 2:163110536 | G/A | 1.15 (1.11-1.2) | 1.27E-11 | 8.9kb 5' of *FAP* |  |
| rs11889341 | 2:191943742 | C/T | 1.70 (1.64-1.75) | 2.48E-75 | *STAT4* Intronic |  |
| rs6736175 (rs16833249) | 2:191946322 | T/C | 1.24 (1.19-1.29) | 9.17E-17 | *STAT4* Intronic |  |
| rs3768792 | 2:213871709 | A/G | 1.24 (1.17-1.31) | 1.21E-13 | *IKZF2* 3'-UTR |  |
| rs9311676 (rs11714389) | 3:58470351 | C/T | 1.17 (1.13-1.22) | 3.06E-14 | *LOC101929223* NC | *ABHD6* |
| rs564799 | 3:159728987 | C/T | 1.14 (1.09-1.18) | 1.54E-09 | *IL12A-AS1* Intronic | *IL12A* |
| rs10028805 | 4:102737250 | G/A | 1.20 (1.15-1.25) | 4.31E-17 | *BANK1* Intronic | *BANK1* |
| rs7726414 (rs17167273) | 5:133431834 | C/T | 1.45 (1.32-1.58) | 4.44E-16 | 19kb 5' of *TCF7* |  |
| rs10036748 | 5:150458146 | C/T | 1.38 (1.32-1.45) | 1.27E-45 | *TNIP1* Intronic |  |
| rs2431697 | 5:159879978 | T/C | 1.26 (1.21-1.31) | 8.01E-28 | 15kb 5' of *hsa-mir-146a* |  |
| rs6568431 | 6:106588806 | C/A | 1.21 (1.15-1.27) | 5.04E-14 | 31kb 3' of *PRDM1* |  |
| rs6932056 | 6:138242437 | T/C | 1.83 (1.65-2.02) | 1.97E-31 | 22kb 3' of *RP11-10J5.1* |  |
| rs849142 | 7:28185891 | T/C | 1.14 (1.1-1.19) | 8.61E-11 | *JAZF1* Intronic |  |
| rs4917014 | 7:50305863 | T/G | 1.18 (1.13-1.24) | 6.39E-14 | 8.5kb 3' of *AC020743.4* |  |
| rs3757387 (rs4728142) | 7:128576086 | T/C | 1.45 (1.4-1.5) | 1.14E-48 | 1.6kb 5' of *IRF5* |  |
| rs35000415 (rs10488631) | 7:128585616 | C/T | 1.83 (1.76-1.9) | 1.20E-60 | *IRF5* Intronic | *IRF5, TNPO3* |
| rs2736340 | 8:11343973 | C/T | 1.29 (1.22-1.37) | 6.28E-20 | 7.5kb 5' of *BLK* | *BLK* |
| rs2663052 | 10:50069395 | G/A | 1.16 (1.1-1.22) | 5.25E-09 | *WDFY4* Intronic | *WDFY4* |
| rs4948496 | 10:63805617 | T/C | 1.14 (1.1-1.19) | 1.04E-10 | *ARID5B* Intronic |  |
| rs12802200 (rs2396545) | 11:566936 | C/A | 1.23 (1.15-1.31) | 8.81E-10 | *MIR210HG* NC |  |
| rs2732549 | 11:35088399 | A/G | 1.24 (1.19-1.29) | 1.20E-23 | 46kb 3' of *PDHX* |  |
| rs3794060 | 11:71187679 | T/C | 1.23 (1.18-1.29) | 1.32E-20 | *NADSYN1* Intronic |  |
| rs7941765 (rs6590343) | 11:128499000 | C/T | 1.14 (1.1-1.19) | 1.35E-10 | 547bp 3' of *RP11-744N12.3* |  |
| rs10774625 | 12:111910219 | A/G | 1.13 (1.08-1.18) | 4.09E-09 | *ATXN2* Intronic |  |
| rs1059312 | 12:129278864 | A/G | 1.17 (1.12-1.21) | 1.48E-13 | *SLC15A4* Synonymous |  |
| rs4902562 | 14:68731458 | G/A | 1.14 (1.09-1.19) | 6.15E-10 | *RAD51B* Intronic |  |
| rs2289583 | 15:75311036 | C/A | 1.19 (1.14-1.24) | 6.22E-15 | *SCAMP5* Intronic | *CSK, ULK3, MPI* |
| rs9652601 | 16:11174365 | G/A | 1.21 (1.15-1.26) | 7.42E-17 | *CLEC16A* Intronic |  |
| rs34572943 (rs9936831) | 16:31272353 | G/A | 1.71 (1.61-1.81) | 3.39E-76 | *ITGAM* Intronic |  |
| rs11644034 | 16:85972612 | G/A | 1.25 (1.19-1.32) | 9.58E-18 | 16kb 3' of *IRF8* |  |
| rs2286672 | 17:4712617 | C/T | 1.25 (1.16-1.35) | 2.93E-09 | *PLD2* Missense | *RNF167* |
| rs2941509 | 17:37921194 | C/T | 1.35 (1.22-1.49) | 7.98E-09 | *IKZF3* 3'-UTR |  |
| rs2304256 | 19:10475652 | C/A | 1.24 (1.17-1.31) | 3.50E-13 | *TYK2* Missense |  |
| rs7444 | 22:21976934 | T/C | 1.27 (1.21-1.33) | 1.84E-22 | *UBE2L3* 3'-UTR | *UBE2L3* |
| SLE GWAS SNPs taken forward for *cis*-eQTL association analysis. Results from post-replication meta-analysis as described (57). SNP with the lowest P-value post meta-analysis or the SNP with the greatest evidence of a missense effect as defined by a Bayes Factor reported. Autosomal, non-MHC SNPs with MAF > 0.05 included in analysis only (39 total). Risk alleles are highlighted in bold type – minor allele on right. Functional annotation from HaploReg v4.0 using GENCODE genes v10. Stated eGenes detected from microarray studies in LCLs listed as described (57). |  |  |  |  |  |  |

**Fig. 1:**
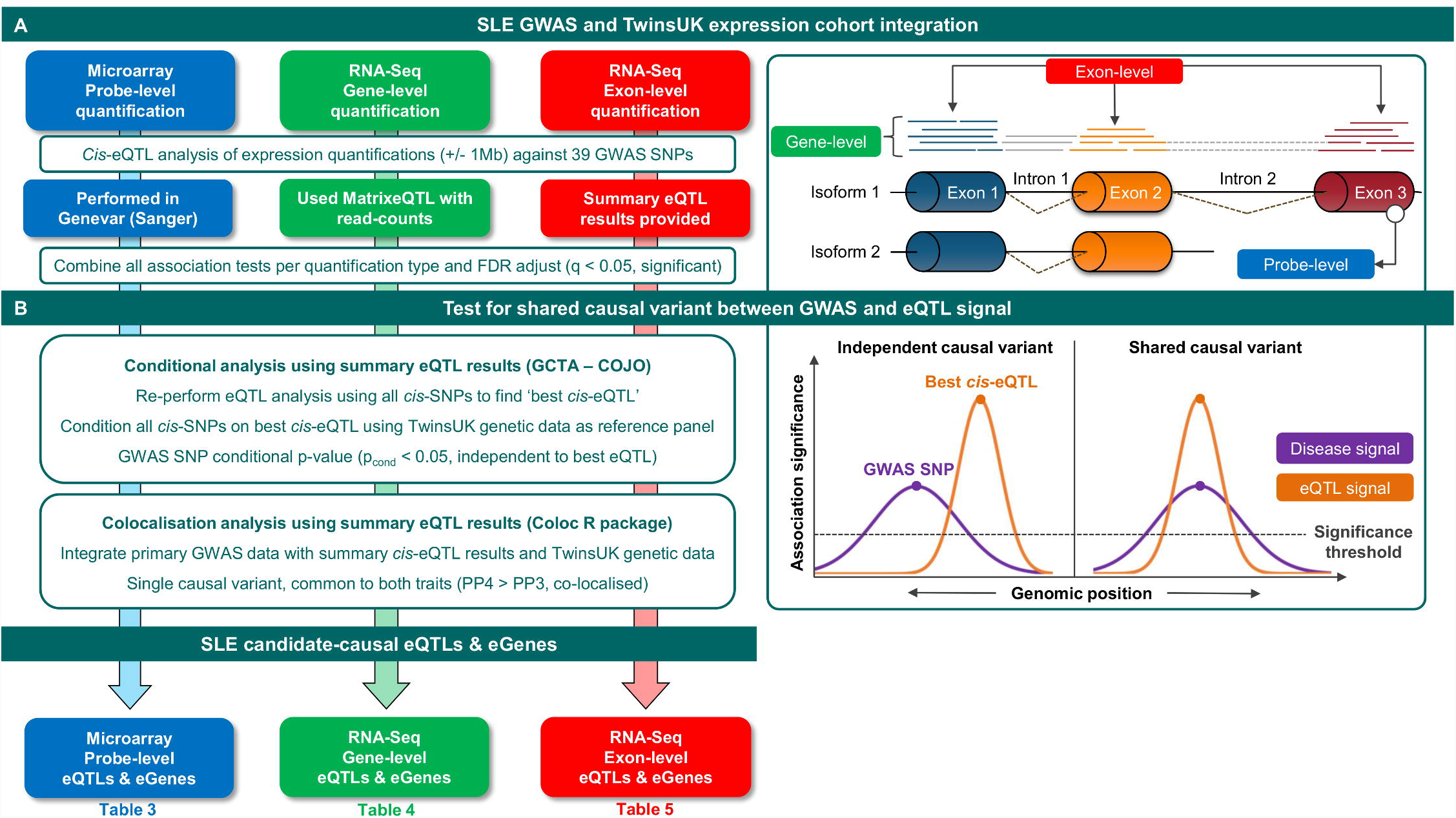
Two-stage *cis*-eQTL annotation pipeline for SLE susceptibility loci. SLE susceptibility variants (**Table 1**) were annotated using residualized expression or summary-level eQTL statistics from three expression datasets: microarray probe-level expression data, and both gene-level and exon-level RNA-Seq quantifications. Each expression dataset was generated from LCLs from individuals of the TwinsUK cohort. **A**) We undertook *cis*-eQTL analysis of +/-1Mb intervals around each SNP and associations with q<0.05 after FDR adjustment were taken forward. **B**) Summary-level data from significant *cis*-eQTLs were tested for evidence of a shared causal variant using firstly conditional analysis using the TwinsUK genetic data as a reference panel, then colocalisation analysis to test for a single causal variant common to both traits. Associations passing these thresholds (described fully in methods) were classified as candidate-causal eQTLs and eGenes. Summary results per quantification type for significant and candidate-causal associations are shown in **Table 3** for microarray, **Table 4** for RNA-Seq (gene-level), and **Table 5** for RNA-Seq (exon-level). Full summary results are available in **Table S1**, **S2**, and **S3** respectively.

**Table 2.**
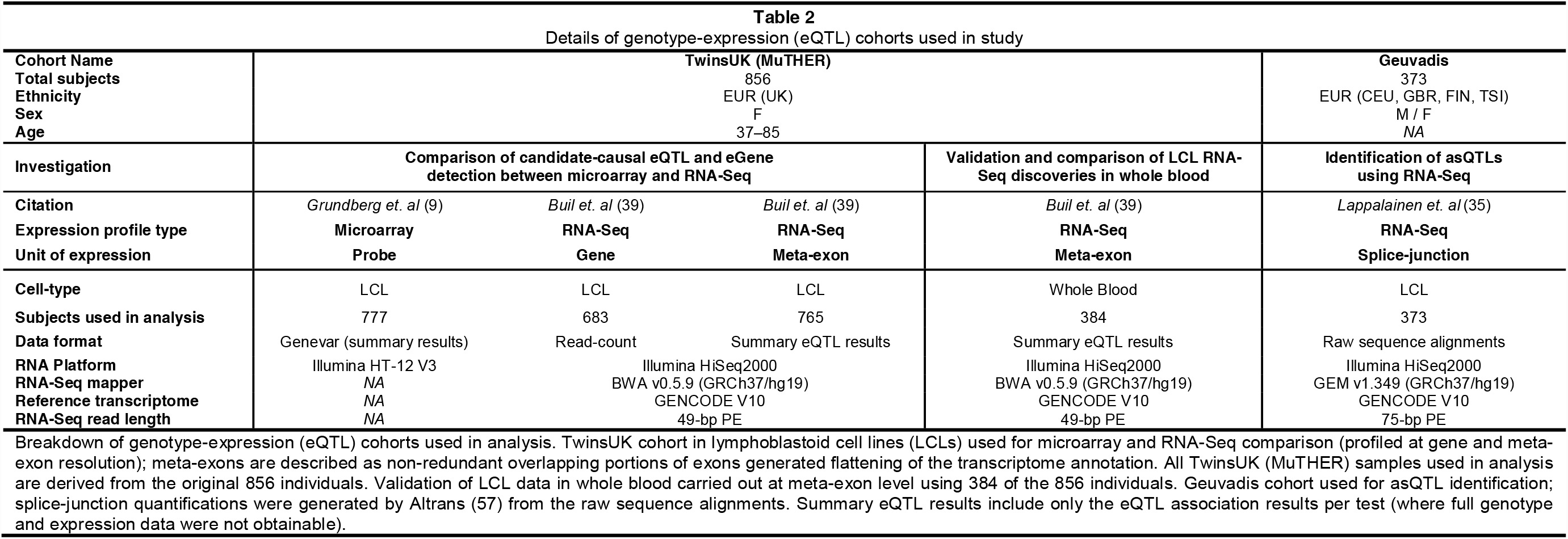
Details of genotype-expression (eQTL) cohorts used in study

Full results of the conditional and colocalisation analysis for each significant association are presented in **S1 Table**, **S2 Table**, and **S3 Table** for microarray, RNA-Seq (gene-level), and RNA-Seq (exon-level), respectively. Statistically significant SLE-associated *cis*-eQTLs showing evidence of a shared causal variant or in very strong LD and close ranking between the disease and *cis*-eQTL signal following conditional and colocalisation analyses were classified as SLE candidate-causal eQTLs as stated in Methods. SLE candidate-causal eGenes were defined as genes whose expression is modulated by the eQTL. The final column of **S1-S3 Tables** indicates whether each GWAS SNP is deemed to be candidate-causal. These SLE candidate-causal eQTLs and eGenes are presented in separate tables contingent on the dataset from which they were generated: results from microarray assessment are listed in **Table 3**, from RNA-Seq (gene-level) in **Table 4**, and from RNA-Seq (exon-level) in **Table 5**. Effect sizes are with respect to the minor allele; risk alleles are highlighted in **Table 1**. Overall, exon-level analysis was the most effective quantification type for the discovery of eQTLs and eGenes compared with gene-level RNA-Seq or microarray analysis following an FDR cut-off of q < 0.05 and conditional and colocalisation thresholding as described.

**Table 3.** Candidate-causal eQTLs and associated eGenes detected using microarray (probe-level)

| Risk Locus | GWAS SNP | eGene | Probe ID | $\beta$ | P-Value | FDR (q) |
| --- | --- | --- | --- | --- | --- | --- |
| 1p13.2 | rs2476601 | *BCL2L15* | ILMN_1655722 | -0.075 | 8.26E-04 | 2.73E-02 |
| 3q25.33 | rs564799 | *IL12A* | ILMN_1671353 | 0.113 | 4.91E-13 | 4.86E-11 |
| 3p14.3 | rs11714389 | *ABHD6* | ILMN_1706344 | -0.130 | 1.41E-13 | 1.63E-11 |
|  |  | *PDHB* | ILMN_1739274 | -0.055 | 2.55E-05 | 1.26E-03 |
| 4q24 | rs10028805 | *BANK1* | ILMN_1661646 | 0.195 | 7.98E-13 | 6.91E-11 |
| 8p23.1 | rs2736340 | *BLK* | ILMN_1668277 | -0.407 | 7.69E-25 | 1.07E-22 |
|  |  | *FAM167A* | ILMN_1687213 | 0.412 | 1.48E-43 | 1.03E-40 |
| 15q24.2 | rs2289583 | *CSK* | ILMN_1754121 | -0.059 | 9.56E-07 | 5.10E-05 |
|  |  | *MPI* | ILMN_1761262 | 0.058 | 1.45E-04 | 5.58E-03 |
|  |  | *ULK3* | ILMN_1679495 | -0.071 | 1.78E-09 | 1.12E-07 |
| 17p13.2 | rs2286672 | *INCA1* | ILMN_1704380 | 0.035 | 7.69E-04 | 2.66E-02 |
| 22q11.21 | rs7444 | *UBE2L3* | ILMN_1677877 | -0.183 | 4.97E-25 | 8.61E-23 |

GWAS SNPs deemed to be candidate-causal eQTLs using microarray expression data profiled from 777 individuals of the TwinsUK cohort in lymphoblastoid cell lines. 768 probes, corresponding to 559 genes, were tested against in *cis* to the 39 GWAS SNPs.

**Table 4.** Candidate-causal eQTLs and associated eGenes detected using RNA-Seq (Gene-level)

|  |  |  |  |  |  |
| --- | --- | --- | --- | --- | --- |
| Risk Locus | GWAS SNP | eGene | $\beta$ | P-Value | FDR (q) |
| 1p13.2 | rs2476601 | *DCLRE1B* | -0.330 | 8.41E-04 | 1.91E-02 |
| 3q25.33 | rs564799 | *IL12A* | 0.372 | 1.07E-11 | 9.70E-10 |
| 3p14.3 | rs11714389 | *ABHD6* | 0.320 | 7.00E-09 | 2.28E-07 |
|  |  | *RPP14* | -0.285 | 4.15E-07 | 1.64E-05 |
| 4q24 | rs10028805 | *BANK1* | -0.316 | 2.37E-08 | 1.29E-06 |
| 5q31.1 | rs17167273 | *SKP1* | -0.564 | 1.57E-05 | 5.06E-04 |
|  |  | *TCF7* | -0.534 | 2.89E-05 | 8.27E-04 |
| 5q33.3 | rs2431697 | *MIR146A* | 0.256 | 1.52E-06 | 6.89E-05 |
| 8p23.1 | rs2736340 | *FAM167A* | 0.668 | 1.40E-33 | 7.62E-31 |
|  |  | *BLK* | -0.596 | 3.13E-28 | 8.51E-26 |
|  |  | *RP11-148O21.2* | -0.591 | 1.11E-26 | 2.01E-24 |
| 11p15.5 | rs2396545 | *HRAS* | -0.248 | 1.56E-04 | 4.04E-03 |
| 11q13.4 | rs3794060 | *NADSYN1* | -0.620 | 6.43E-24 | 8.74E-22 |
|  |  | *DHCR7* | -0.200 | 1.70E-03 | 3.43E-02 |
| 15q24.2 | rs2289583 | *ULK3* | 0.303 | 3.62E-07 | 1.79E-05 |
|  |  | *UBE2Q2* | 0.249 | 5.66E-05 | 1.54E-03 |
|  |  | *FAM219B* | -0.198 | 1.18E-03 | 2.57E-02 |
|  |  | *CSK* | 0.194 | 1.64E-03 | 3.43E-02 |
| 22q11.21 | rs7444 | *UBE2L3* | 0.345 | 2.28E-06 | 9.54E-05 |
| GWAS SNPs deemed to be candidate causal eQTLs using RNA-Seq expression data profiled from 683 individuals of the TwinsUK cohort in lymphoblastoid cell lines at gene-level resolution. 520 genes were tested against in *cis* to the 39 GWAS SNPs. |  |  |  |  |  |

**Table 5.**
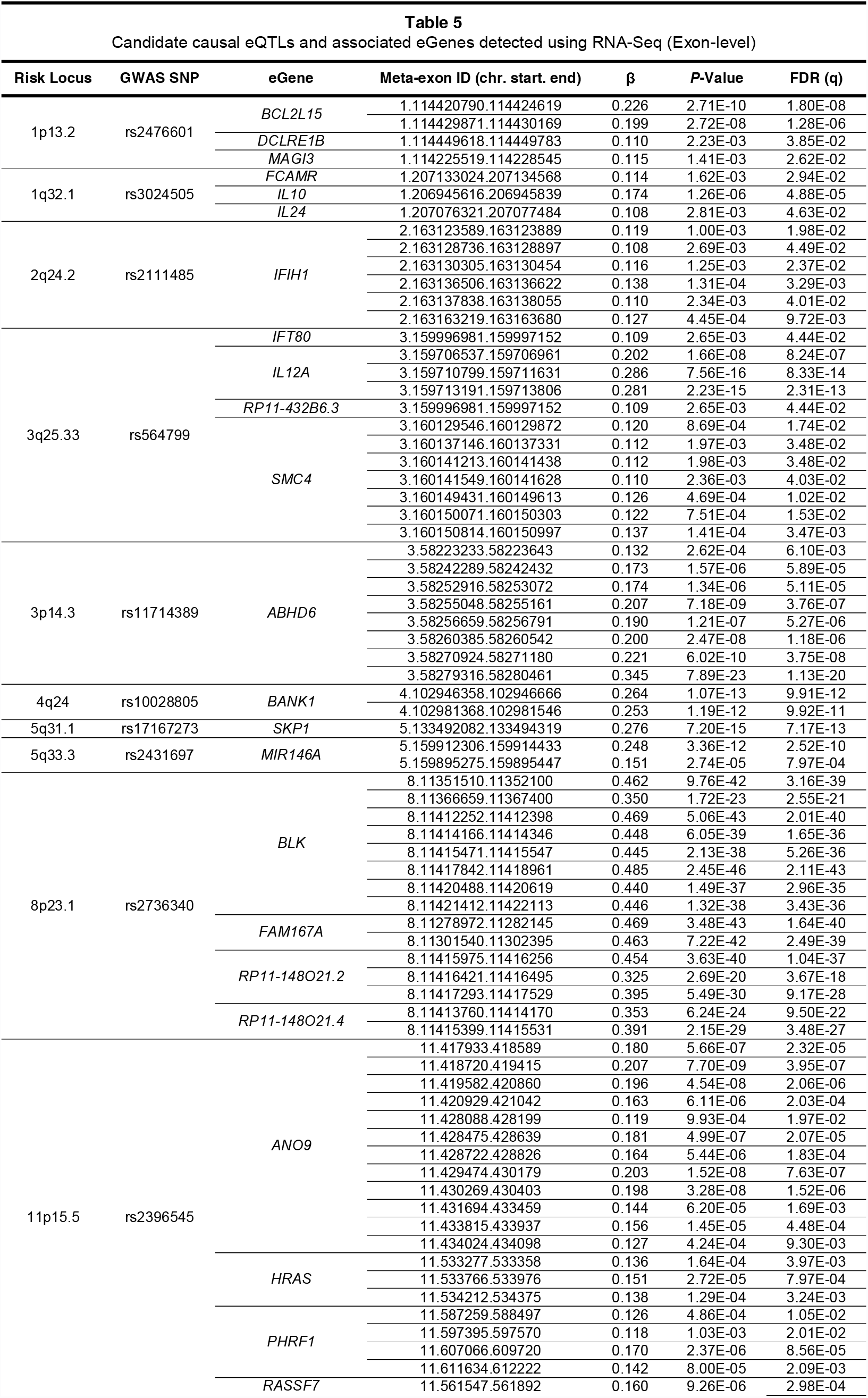
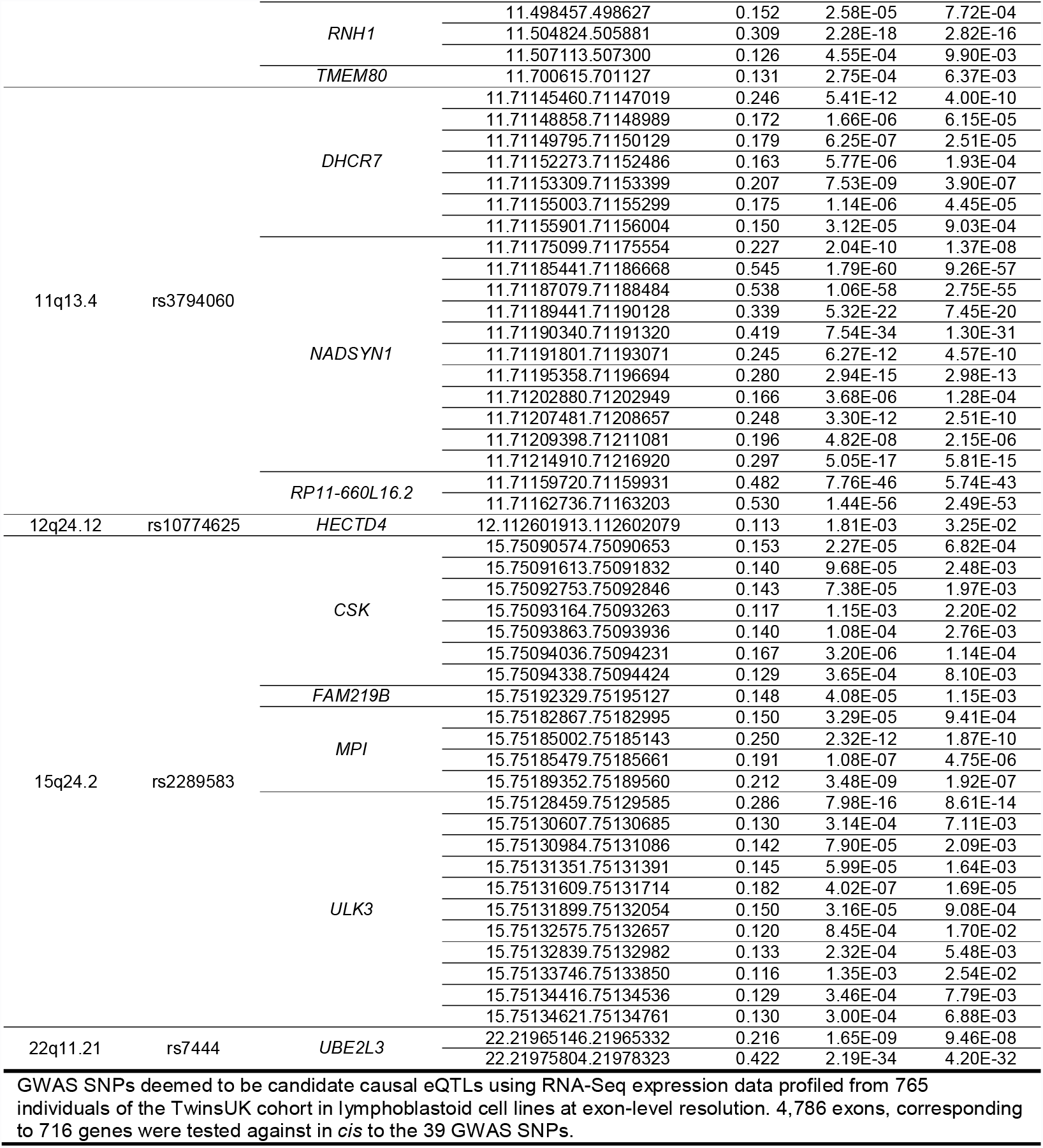
Candidate causal eQTLs and associated eGenes detected using RNA-Seq (Exon-level)

### RNA-Seq better annotates SLE risk loci by revealing known and novel candidate genes

**Fig. 2** illustrates the clear improvement of RNA-Seq relative to microarray in the discovery of candidate-causal eQTLs and their corresponding eGenes when annotating complex-disease susceptibility loci. With RNA-Seq, we replicated known SLE associated eQTLs and eGenes that were previously discovered using microarray. These included rs564799 for *IL12A*, rs2736340 for *BLK*, rs9311676 *ABHD6*, and rs2289583 for *ULK*, *CSK*, and *MPI* (**Tables 4**, and **5**; **Table 1** for previously reported associations in LCLs). Several eQTLs for eGenes that have been extensively studied in terms of their role in SLE pathogenesis were consistent across all three platforms: for example, rs10028805 for *BANK1* (57) and rs7444 for *UBE2L3* (58). **Fig. 2** also shows how exon-level RNA-Seq analysis led to the greatest frequency of candidate-causal eQTLs and eGenes than with either gene-level RNA-Seq or microarray (**S1 Fig.** and **S2 Fig.**). A total of 14 eQTLs modulating expression of 34 eGenes were detected using exon-level RNA-Seq contrasted to 11 eQTLs and 19 eGenes at gene-level RNA-Seq and only 8 eQTLs with 12 eGenes identified using microarray (**S1 Fig.** and **S2 Fig.**). Only one eQTL (rs2286672) and two eGenes (*PDHB*, *INCA1*) were found by microarray exclusively. These associations were either not significant post multiple testing using either RNA-Seq method, or were not deemed candidate-causal (**S1 Table S1** and **S3 Table**). In total 8 eQTLs regulating expression of 27 eGenes were detected using RNA-Seq but missed using microarray (**S3 Fig. S3** and **S4 Fig.**). Interestingly, exon-level analysis led to the greatest frequency of non-candidate-causal associations. Only 14 of the 34 significant eQTLs, q < 0.05, showed evidence of a shared causal variant post conditional and colocalisation testing (**S1 Fig.** and **S2 Fig.**).

**Fig. 2:**
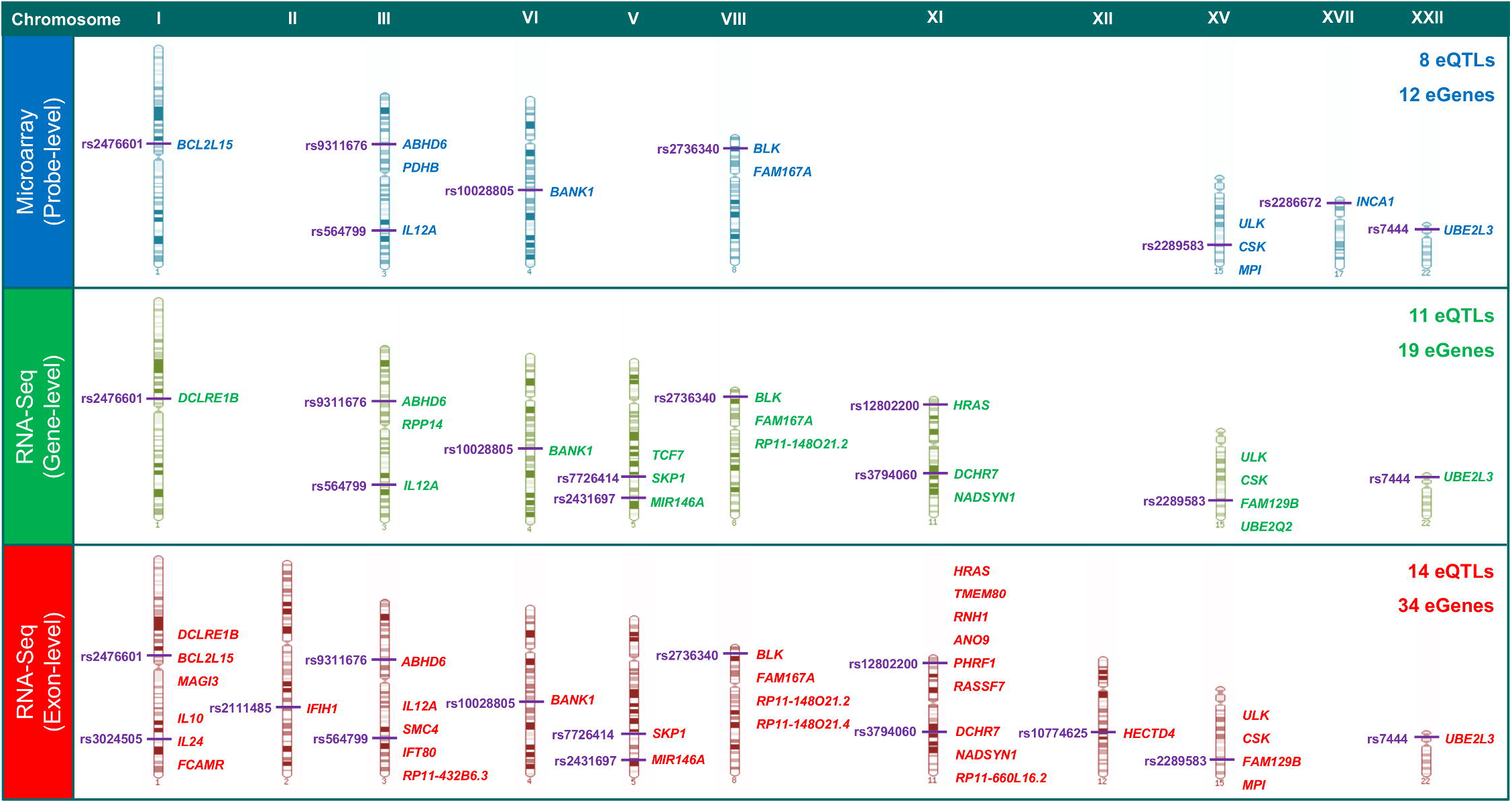
Candidate-causal eQTLs and eGenes detected across quantification types. Candidate-causal eQTLs and eGenes detected using microarray (probe-level), RNA-Seq (gene-level), and RNA-Seq (exon-level) quantifications from the TwinsUK datasets are represented in their genomic context (**Tables 3**, **4**, and **5** respectively). Only chromosomes harbouring one or more candidate-causal eQTL or eGene per quantification type are shown.

Several eGenes known to be involved in the pathogenesis of SLE were identified using RNA-Seq exclusively and not reported in previous microarray-based eQTL studies in LCLs (**Table 1**). These include *IL10*, *IFIH1*, and the microRNA *MIR146A*. We believe a handful of other eGenes unique to RNA-Seq to be novel SLE candidate genes. *NADSYN1* (NAD Synthetase), *HECTD4* (HECT Domain E3 Ubiquitin Protein Ligase 4), *SKP1* (S-Phase Kinase-Associated Protein 1) and *TCF7* (T-cell specific Transcription Factor 7) are examples of novel eGenes.

### RNA-Seq eQTL analysis reveals eQTLs regulating multiple eGenes

**Fig. 2** also illustrates that many of the eQTLs discovered using RNA-Seq regulate multiple eGenes. Exon-level analysis generated the greatest ratio (2.42) of candidate-causal eGenes to eQTLs (**S5 Fig.**); suggesting that disease-associated haplotypes may be more functionally potent and harbour multiple gene regulatory effects than previously thought.

One example of this effect is at the *TCF7*-*SKP1* locus where the disease-association signal encompasses both genes (**S6 Fig.**). Both *TCF7* and *SKP1* were classified as candidate-causal against the GWAS SNP rs7726414 using gene-level RNA-Seq (**Table 5**), and *SKP1* but not *TCF7* at exon-level (**Table 4**). Interestingly, a missense variant of *TCF7* has been implicated in Type 1 Diabetes risk (59), but there is only weak LD (r^2^<0.4) between this missense variant and rs7726414 (19 kb upstream of *TCF7*) or any protein-coding variants of *TCF7*, suggesting that in SLE the causal mechanism may be dysregulated gene expression of *TCF7* rather than a missense change *per se*. *SKP1*, part of the ubiquitin ligase complex, is thought to stabilize the conformation of E3 ligases and its expression has been shown to be upregulated in lung-cancer (60). Neither of these effects were present in the microarray data as there is another more significant *cis*-eQTL, rs17167273, for the *TCF7* probe (ILMN_1676470), which is not correlated with the rs7726414[T] risk variant (r^2^:0.02, **S1 Table**). In addition, rs7726414[T] was not a significant eQTL for the *SKP1* microarray probe (ILMN_1790710) (*P*=0.16). Our RNA-Seq eQTL analyses indicated that rs7726414[T] represents a novel eQTL for dysregulated gene expression of *SKP1* in SLE (**S6 Fig.**). We believe both *TCF7* and *SKP1* to be highly plausible candidate genes as the literature suggests that knockdown of *TCF7* results in impaired stem cell potency and gene expression regulation of CD34+ cells (61), whilst the mouse knockout of *SKP1* develop highly penetrant T-cell lymphomas (62).

Other examples where either gene- or exon-level RNA-Seq analysis identified multiple eGenes are at rs3024505 (*IL10*, *IL24*, and *FCAMR*), rs12802200 and rs3794060 (**Fig. 2**). In the immune-gene concentrated 11p15.5 region (**S7 Fig.**), the GWAS SNP rs12802200, is an eQTL for six eGenes, (*HRAS, TMEM80*, *RNH1*, *ANO9*, *PHRF1* and *RASSF7*; **Tables 4 and 5**) following exon- and/or gene-level RNA-Seq analysis, supporting our recent observations of rs12802200 being a *cis*-eQTL for six genes across multiple immune cell-types at this locus (63) and earlier reports demonstrating that rs12802200 was an eQTL for *TMEM80* (whole blood(64)) and for both *RNH1* and *TMEM80* (LCLs (35)). 11p15.5 is also an important regulatory region in non-immune types, because a total of eight eGenes in this region have been identified in multiple non-immune cell types (65), four of which (*HRAS*, *TMEM80*, *RNH1*, *ANO9*) were captured by our RNA-Seq analysis in LCLs. Since SNPs within this susceptibility locus have also been previously shown to be correlated with increased autoantibody production and interferon-α activity in sufferers of SLE (66), further investigation will be required to elucidate a potential mechanism by which dysregulation of gene expression may contribute to these effects. In this gene-dense region (**S7 Fig.**), none of the six candidate-causal eGenes had an annotated microarray probe that passed quality control.

In the 11q13.4 region, the intronic GWAS SNP rs3794060[C], was classified using gene-level quantification as being a candidate-causal eQTL for both *NADSYN1* (NAD Synthetase 1) and *DHCR7* (7-Dehydrocholesterol Reductase) (**Table 4, Fig. 3A** and **S8 Fig.**). At exon-level resolution, *NADSYN1* was also deemed candidate-causal, as well as the non-coding RNA *RP11-660L16.2* (**Table 5**; described in following section). The risk variant rs3794060[C] has been correlated with reduced circulating 25-hydroxy vitamin D concentrations (67) with reduced vitamin D levels being recently associated with increased disease activity of SLE (68). Interestingly, although a mouse knockout of *DHCR7* showed reduced serum and tissue cholesterol levels (69), no autoimmune phenotype has been described at this locus. Given the role oxidative stress plays in promoting inflammation and triggering autoimmunity through tissue damage (70), it will be interesting to elucidate the role that dysregulation of *NADSYN1* expression plays in this process. Visualization of exon-level association data of *NADSYN1* against rs3794060 suggests a potential splicing mechanism affecting meta-exons 11 and 12 (*P*=1.79^E-60^, 1.06^E-58^ respectively; **Fig. 3B**). The probe for *DHCR7* (ILMN_1815626) showed a weak, but not statistically significant, association with rs3794060 (*P*=2.9x10^-03^), whilst *NADSYN1* had no annotated probe in the microarray dataset.

**Fig. 3:**
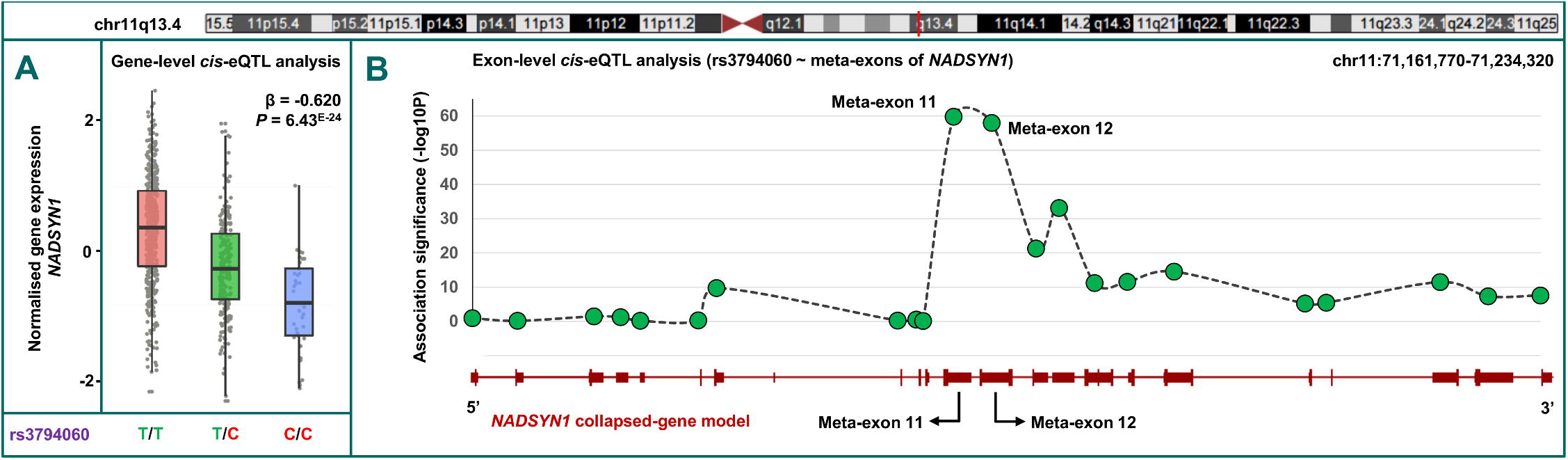
RNA-Seq gene-level and exon-level analysis implicate *NADSYN1* as a novel candidate-causal eGene. **A**) *Cis*-eQTL analysis of GWAS SNP rs3794060 reveals the risk variant [C] leads to substantial down-regulation at the gene-level of novel susceptibility gene *NADSYN1* in the TwinsUK cohort that was not detected using microarray. **B**) The same analysis using exon-level quantification leads to the inference of the gene-level effect being driven by considerable expression disruption of two meta-exons of *NADSYN1* (meta-exon 11 and meta-exon 12). Association *P*-values of rs3794060 against exon quantifications are plotted with reference to the specific exon in the collapsed-gene model of *NADSYN1* (all annotated transcripts combined).

### RNA-Seq uncovers the role of non-coding RNA modulation at SLE susceptibility loci

Quantification of non-coding polyadenylated RNAs in the TwinsUK LCL cohort through RNA-Seq revealed three candidate-causal eQTLs influencing the expression of four non-coding eGenes (**Table 6**); none of which were captured using microarray.

**Table 6.**
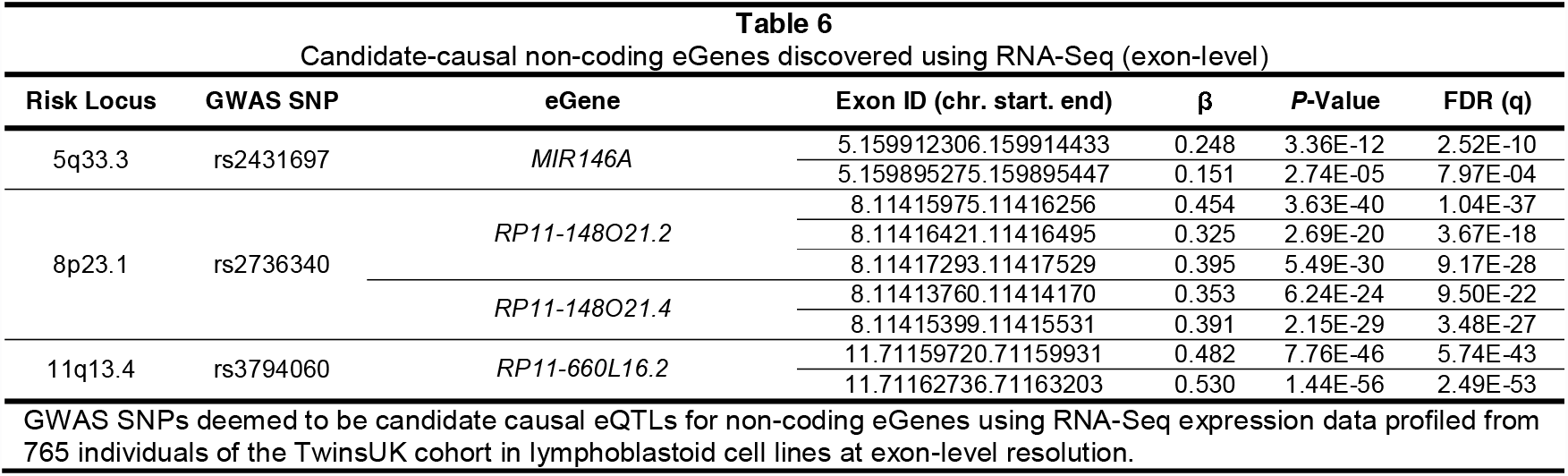
Candidate-causal non-coding eGenes discovered using RNA-Seq (exon-level)

We validated the effect at rs2431697 (5q33.3) where it is documented that the protective minor allele [C] is associated with expression upregulation of the miRNA *MIR146A*, a negative regulator of the type I Interferon pathway (71) (**Fig. 4A**). The best eQTL for *MIR146A* at gene-level was rs2431697 (*P*=1.5×10^-06^) and also at exon level for both of its exons (*P*=3.4×10^-12^ and 1.2×10^-04^). The decrease in rs2431697[T]-dependent expression of *MIR146A* reported in peripheral blood leukocytes of SLE patients disrupts binding of transcription factor Ets-1, uncouples of the type-1 IFN response (71), thereby increasing the inflammatory response.

**Fig. 4:**
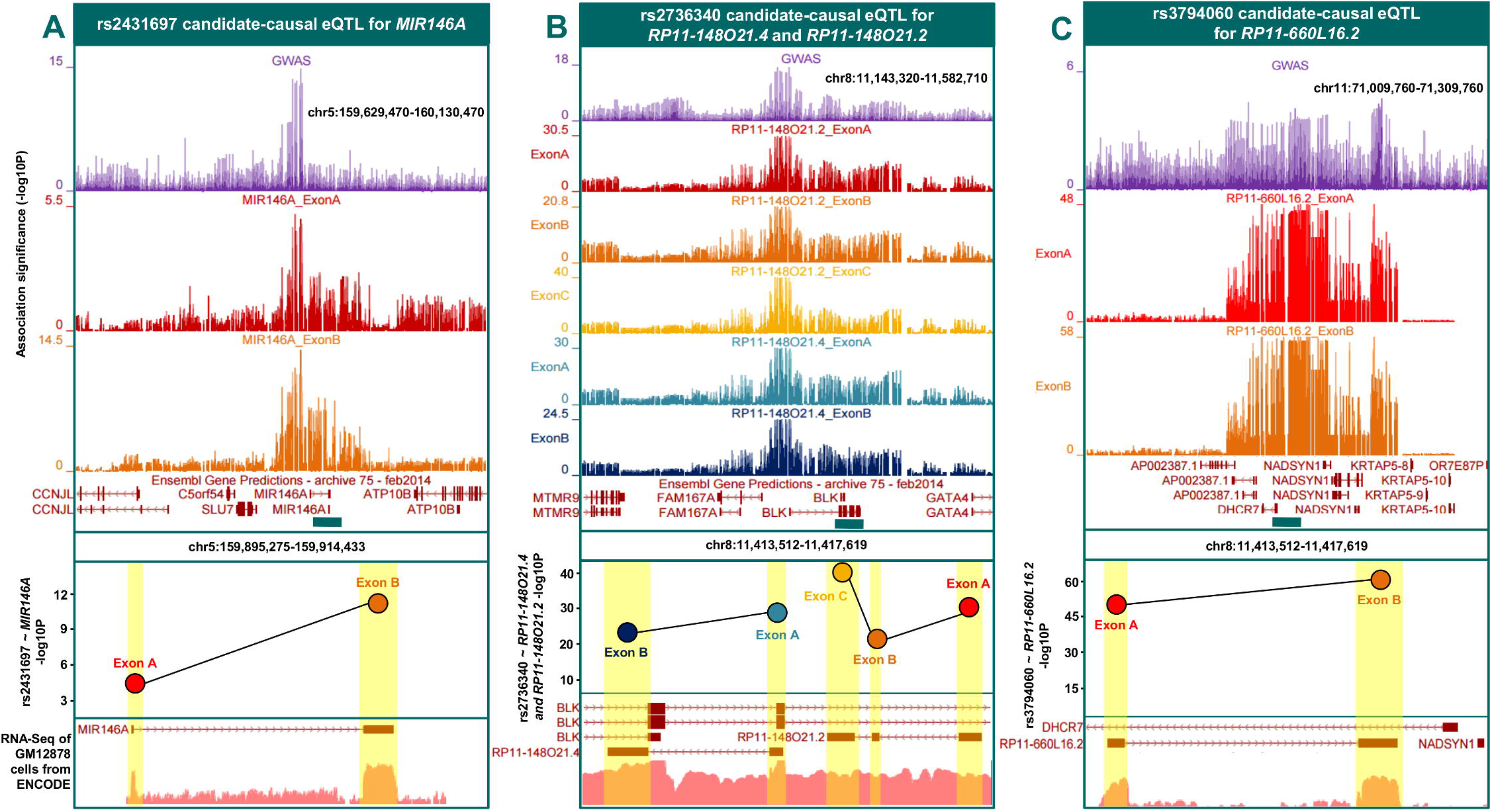
Non-coding candidate-causal eGenes detected using exon-level RNA-Seq. The three figures denote the eQTLs and corresponding non-coding eGenes identified from *cis*-eQTL analysis of GWAS SNPs against TwinsUK exon-level quantifications in LCLs. The top panels display the signal from the GWAS association plotted as –log_10_ (*P*), below which the exon-level eQTL *P*-values for the effects showing colocalisation with the GWAS signal are illustrated. The bar across the middle of the figure denotes the boundaries of the eQTL, below which there is a panel showing the association *P*-value of the GWAS SNP against the candidate-causal non-coding exons. The bottom panel shows LCL RNA-Seq alignments from ENCODE to show that the regions containing the candidate-causal eQTLs are expressed. **A**) GWAS SNP rs2431697 is a candidate-causal eQTL for non-coding eGene *MIR146A*. **B**) GWAS SNP rs2736340 is a candidate-causal eQTL for non-coding eGenes *RP11-148O21.4* and *RP11-148O21.2*. **C**) GWAS SNP rs3794060 is a candidate-causal eQTL for non-coding eGene *RP11-660L16.2*.

**Fig. 4B** shows GWAS variant rs2736340, and other SNPs in tight LD (r^2^>0.8), within the *FAM167A*-*BLK* (8p23.1) bi-directional promoter region. In this study we detected eQTLs at the known eGenes *BLK* (B lymphocyte kinase) and *FAM167A* (Family with Sequence Similarity 167, Member A) by all quantification methods (**S9 Fig.**). The rs2736340[T] risk allele causes decreased expression of *BLK* (*P*=3.2×10^-28^) and increased expression of *FAM167A* (*P*=1.4×10^-33^) at RNA-Seq gene-level (**Table 4**). This effect has been previously described in microarray studies (72) - with reduced promoter activity of *BLK* leading to altered B-cell development (73). The eQTL lies within a region of the genome subject to multiple regulatory effects, with a 24bp region immediately around the variant containing strong chromatin marks (peak height>60) in LCLs for H3K9me3 (gene repression) and H3K4me3 (low expression when present in the promoter of CpG genes) assayed as part of the ENCODE project (**S9 Fig.**). Interestingly, exon-level RNA-Seq analysis revealed that rs2736340 also appeared to modulate the expression of two non-coding RNAs antisense to the 3′ region of *BLK*. These are: *RP11-148O21.2* and *RP11-148O21.4* (**Fig. 4B**). The SNP, rs2736340, significantly reduced the expression of both *BLK* and three exons of *RP11-148O21.2* and the two exons of *RP11-148O21.4* (**Table 6**). Our analyses indicated that expression disruption of these antisense RNAs caused by SLE risk variants represent a potential novel mechanism at the locus.

There is an rs3794060 allele-dependent expression modulation of both exons of another non-coding eGene, *RP11-660L16.2I* (**Fig. 4C**), which is located in the bi-directional promoter between *DHCR7* and *NADSYN1*. The best eQTL for those exons is highly correlated with the GWAS SNP (rs2282621, r^2^:0.99). Both *DHCR7* and *NADSYN1* are candidate-causal eGenes regulated in the same downward direction at RNA-Seq gene-level with respect to the risk allele rs3794060[T] (*P*=1.7×10^-03^ and *P*=6.4×10^-24^, respectively) (**S8 Fig.** and **Table 4**).

### Confirmation of LCL candidate-causal eQTLs and eGenes using whole-blood RNA-Seq

To validate our LCL findings in a primary tissue-type, we extended our analytical pipeline to include an exon-level RNA-Seq dataset in 384 whole-blood samples from the TwinsUK cohort (**Table 2**). The full results of these analyses are provided in **S4 Table**.

We observed good correlation between LCLs and whole-blood effect-sizes (β) of GWAS SNPs against all matched *cis* exon-level associations (*R*^*2*^=0.74; **S10 Fig.**). Seven of the 39 GWAS SNPs were classified as candidate-causal eQTLs in whole-blood, modifying 19 candidate-causal eGenes (**Table 7**). All seven of the whole-blood eQTLs and 15 of the 19 eGenes were deemed candidate-causal in LCLs, suggesting strong conservation across whole-blood cell types (**S11 Fig.**). The remaining four eGenes specific to whole-blood were: *PXK* (rs9311676); *IRF7* and *TALDO1* (rs12802200); and *SCAMP2* (rs2289583) (**Table 7**). Interestingly, the eQTLs regulating these four eGenes in whole-blood also regulated multiple eGenes in LCLs (**S11 Fig.**), implying that they tag highly regulatory haplotypes that may cause cell-type specific gene-expression disruption across the entire locus (same eQTL regulating different eGenes across different cell-types). Three of the four candidate-causal non-coding eGenes from LCLs were found in whole-blood (*RP11-148O21.2*, *RP11-148O21.4*, and *RP11-660L16.2*.). The LCL-specific *MIR146A* eGene association with GWAS SNP rs2431697 was not deemed to be significant (*P*=0.32), which is likely to be a result of its lymphocyte-specific gene expression profile (71) that is diluted in the heterogeneous population of whole-blood cell-types.

**Table 7.**
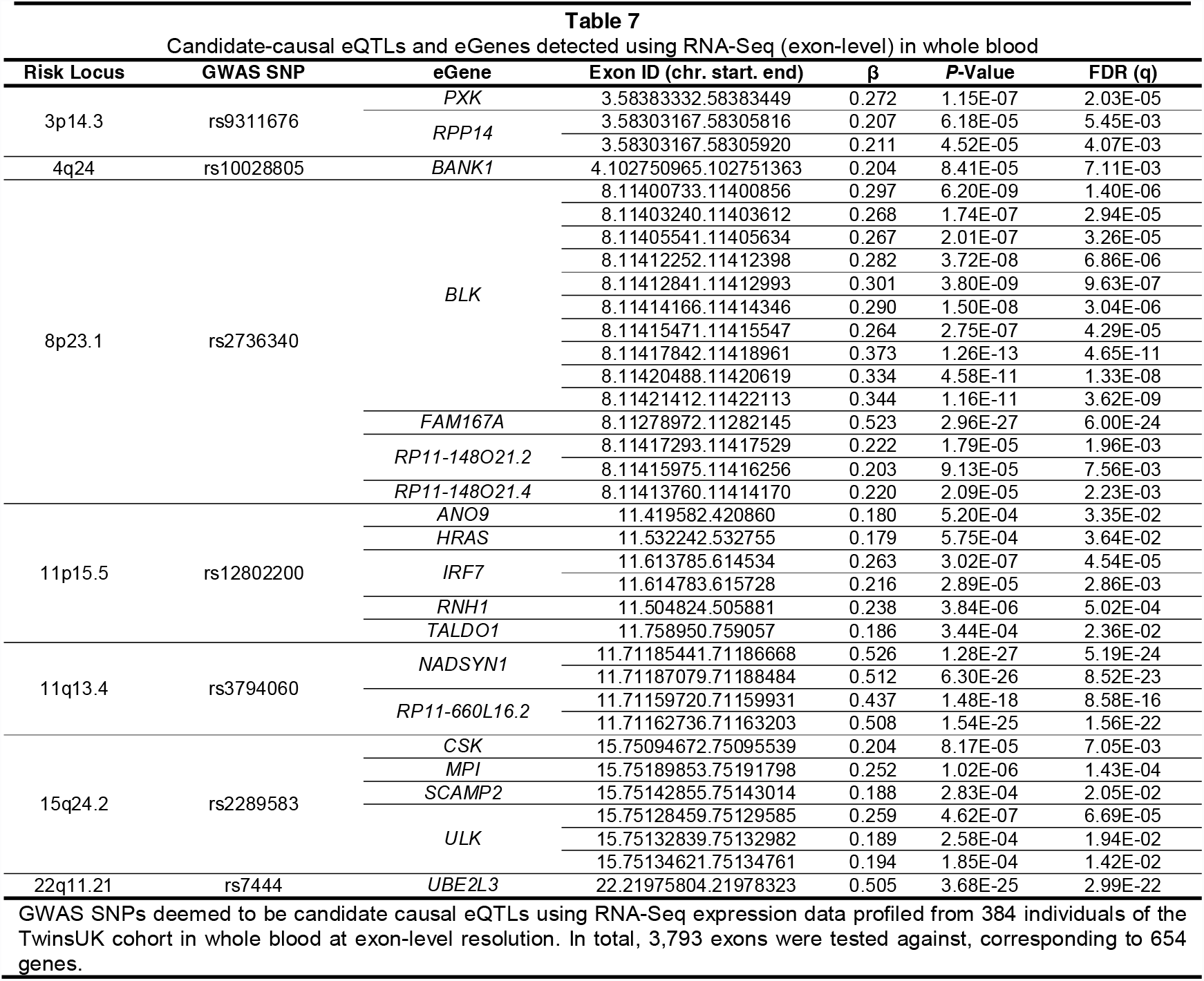
Candidate-causal eQTLs and eGenes detected using RNA-Seq (exon-level) in whole blood

Inspection of specific exons modulated by the GWAS SNPs in each cell-type revealed instances of variability in the genetic control of exon usage between cell-types. A known splicing event in B-cells caused by branch-point SNP rs17266594 results in the loss of exon 2 in susceptibility gene *BANK1* which subsequently leads to B-cell hyper-responsiveness(57) (**S12 Fig.**). In whole-blood the GWAS variant, rs10028805, is associated with altered expression of exon 2 (*P*=8.4×10^-05^), with the best *cis*-eQTL for this effect being in near-perfect LD (rs4411998; r^2^:0.98). Both rs10028805 and rs4411998 are in strong LD with the branch-point SNP (r^2^:0.9). In LCLs however, the best *cis*-eQTL for exon 2, rs4572885 (*P*=9.74 ×10^-23^), has a large effect but is less correlated with the GWAS SNP (r^2^:0.65) and conditional analysis judges the effect of the GWAS SNP to be independent to the best *cis*- eQTL for exon 2 (**S3 Table**). Interestingly, there is low correlation between the branch-point SNP rs17266594 and the best *cis*-eQTL for exon 2 in LCLs (r^2^:0.42); suggesting the regulatory mechanism of exon 2 splicing in *BANK1* may be under two separate genetic influences between the two cell-groups (**S12 Fig.**).

We saw a near identical pattern of differential exon usage within eGene *NADSYN1* between LCLs and whole-blood driven by the GWAS SNP rs37940460 or tightly correlated variants (**S13 Fig.**, **Table 7**). Variation at rs37940460 appeared to drive extensive expression disruption of two meta-exons (11 and 12) of *NADSYN1* located near the centre of the gene (meta-exon 11: LCL *P*=1.79×10^-60^; whole-blood *P*=1.28×10^-27^; meta-exon 12: LCL *P*=1.06×10^-58^; whole-blood *P*=6.30×10^-26^). These two meta-exons were deemed to be candidate-causal for SLE across both cell-types. We believe the meta-exons in the 3′ end of *NADSYN1* that are candidate-causal in LCLs (**Fig. 3**), are not detected in whole-blood may be because of the smaller sample size or the mixed cell-type composition of the whole blood cohort. This novel instance of specific exon expression disruption found in a primary cell-type at *NADSYN1* may help to resolve the functional consequence of this locus.

### Splice-junction quantification reveals asQTLs and additional candidate eGenes

We extended our investigation to determine whether the GWAS SNPs (**Table 1**) had a direct influence on the alternative-splicing of transcripts (alternative-splicing quantitative trait loci; asQTL), and whether expression quantification at this resolution would reveal any additional candidate-genes or potential functional mechanisms. We undertook *cis*-asQTL analysis within a +/-1Mb window around each GWAS SNP against 33,039 splice-junction quantifications, corresponding to 817 genes, in the Geuvadis cohort (**Table 2**). We identified nine asQTLs significantly associated with 62 splice-junctions, corresponding to 10 eGenes (**S5 Table**). After testing for a shared causal variant between the GWAS and asQTL signal, six SLE candidate-causal asQTLs (26 splice-junctions) for seven eGenes remained (**Table 8**). Four eGenes (*TCF7*, *SKP1*, *BLK*, and *NADSYN1*) had been previously associated through either gene-level or exon-level eQTL mapping using the TwinsUK cohort. The remaining three candidate-causal eGenes detected using asQTL mapping (*IKZF2*, *WDFY4*, and *IRF5*), as well as the novel causal mechanism involving *NADSYN1*, are described below.

**Table 8.**
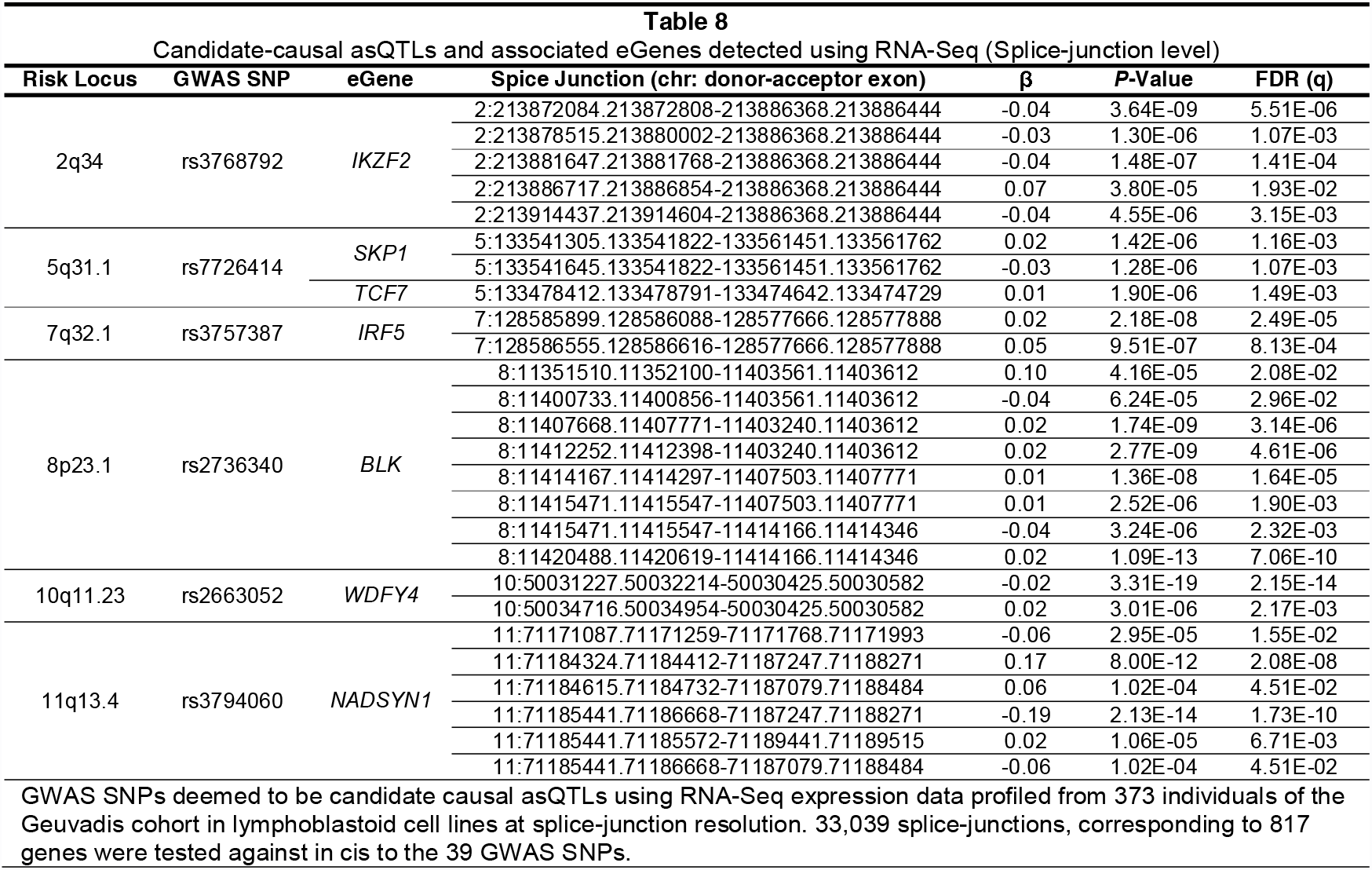
Candidate-causal asQTLs and associated eGenes detected using RNA-Seq (Splice-junction level)

#### IKZF2

*IKZF2* (Ikaros Family Zinc Finger 2) is novel SLE candidate-causal eGene detected only by asQTL analysis. The GWAS association signal around the 3′ end of *IKZF2* tagged by risk variant rs3768792[G] drove an increase in the fraction of splicing between exon 6A and exon 6B (*P*=3.8x10^-05^); a bridge that is unique to the truncated isoform (ENST00000413091, 239 amino-acids) of *IKZF2* (**Fig. 5A**). Interestingly, this isoform possesses a premature termination codon found on exon 6B that is not found on the canonical isoform (ENST00000457361, 526 amino-acids) as in this isoform, exon 6A is spliced to exon 7 (**Fig. 5B**). This effect results in the premature truncation of the full-length protein and the subsequent loss of the two zinc-finger dimerization domains found on exon 8 (**Fig. 5B**). *IKZF2* is a transcription factor thought to play a key role in T-reg stabilisation in the presence of inflammatory responses (74). Since the Ikaros transcription factor family primarily regulate gene expression through homo-/hetero-dimerization and DNA binding/protein-protein interactions, the rs3768792[G] dependent asQTL effect on exon 6A to 6B resulting in less functional *IKZF2* could be highly deleterious. *IKZF2* is known to regulate T-reg associated genes, including *IL-2* and *FoxP3* (75, 76), therefore a decrease in the amount of DNA binding *IKZF2* may result in loss of T-reg stability and a decrease of suppressive capacity with consequential autoimmune sequelae. Interestingly, we identified an additional asQTL variant (rs2291241) in near-perfect LD with the rs3768792 GWAS variant (r^2^:0.99), located 9 bp upstream of exon 6B in truncated isoform ENST00000413091 (**S5 Table, Fig. 5C**). This second asQTL, located within the polypyrimidine tract in the exon 6A/exon 6B intron, is a highly plausible driving variant and may act through promotion of the described splicing event (**Fig. 5C**).

**Fig. 5:**
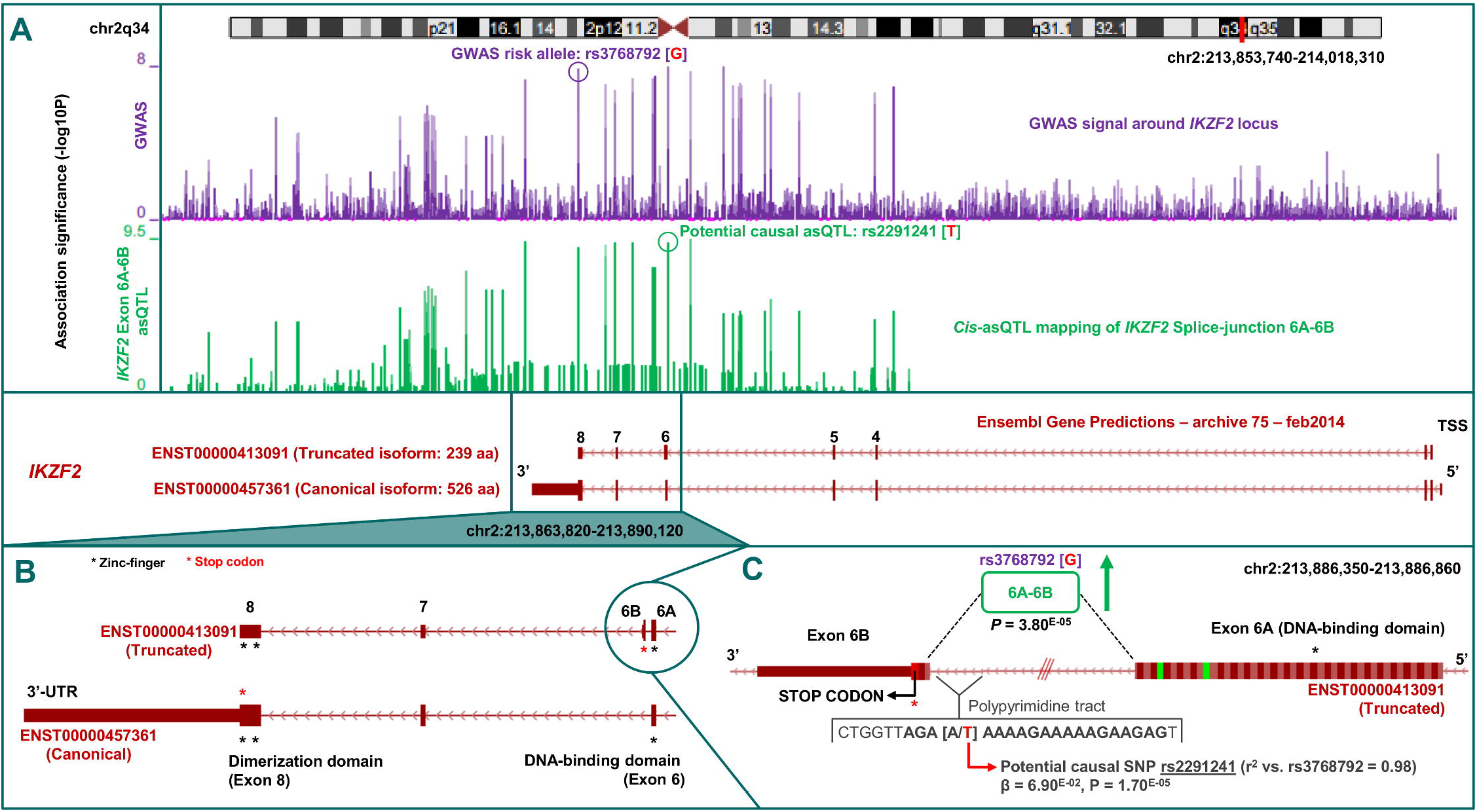
Identification of novel eGene *IKZF2* and potential causal mechanism using RNA-Seq splice-junction quantification. *Cis*-asQTL analysis of GWAS SNP rs3768792 against splice-junction quantifications classified *IKZF2* as a candidate-causal eGene with risk variant [G] causing upregulation of the exon 6A–exon 6B junction that is unique to truncated isoform ENST00000413091. **A**) GWAS association signal across the *IKZF2* locus (chr2q34), tagged by rs3768792 localised in the 3′-UTR of *IKZF2*. *Cis*-asQTL association signal of rs3768792 against splice-junction quantification of exon 6A–exon 6B shows significance and colocalisation with the GWAS signal. **B**) The exon 6A–exon 6B junction is unique to truncated isoform ENST00000413091. Exon 6B harbours a premature stop-codon and therefore is not translated into the full-length protein that contains the dimerization domains in exon 8. **C**) Close-up of the exon 6A–exon 6B junction and association (*P*=3.80E^-05^) with GWAS SNP rs3768792. A potential causal asQTL in near-perfect LD was identified that is located within the polypyrimidine tract of the junction and may induce splicing (rs2291241, *P*=1.70^E-05^).

#### WDFY4

We also discovered a novel putative SLE-associated splicing mechanism involving *WDFY4* (WDFY Family Member 4), a gene belonging to a family thought to function as master conductors of aggregate clearance by autophagy (77). Risk variant rs2263052[G] or correlated SNPs (**Fig. 6A**) greatly increased the fraction of link-counts between exon 34A and exon 34B (*P*=3.3×10^-19^) which are unique to the truncated isoform ENST00000374161 (**Fig. 6B**). This truncated isoform (552 amino-acids) lacks the two WD40 domains found in the full length isoform (ENST00000325239, 3184 amino-acids) that are essential to enzymatic activity (77). There is a consequential decrease in the fraction of link-counts between exon 34A and exon 35 (*P*=3.0×10^-06^) that are unique to the canonical isoform of *WDFY4* (**Fig. 6B**). Interestingly, a known missense variant found in exon 31 of *WDFY4* (**Fig. 6B**), rs7097397 (Arg1816Gln), in strong LD (r^2^:0.7) with rs2263052, has also been implicated in SLE through GWAS (78). SIFT and PolyPhen predict the amino-acid substitution to be tolerated (0.38) or benign (0.11) respectively (79); thereby suggesting the risk haplotype may harbour two functional mechanisms influencing *WDFY4* (amino-acid change and upregulation of a shorter isoform) that are both involved in SLE pathogenesis.

**Fig. 6:**
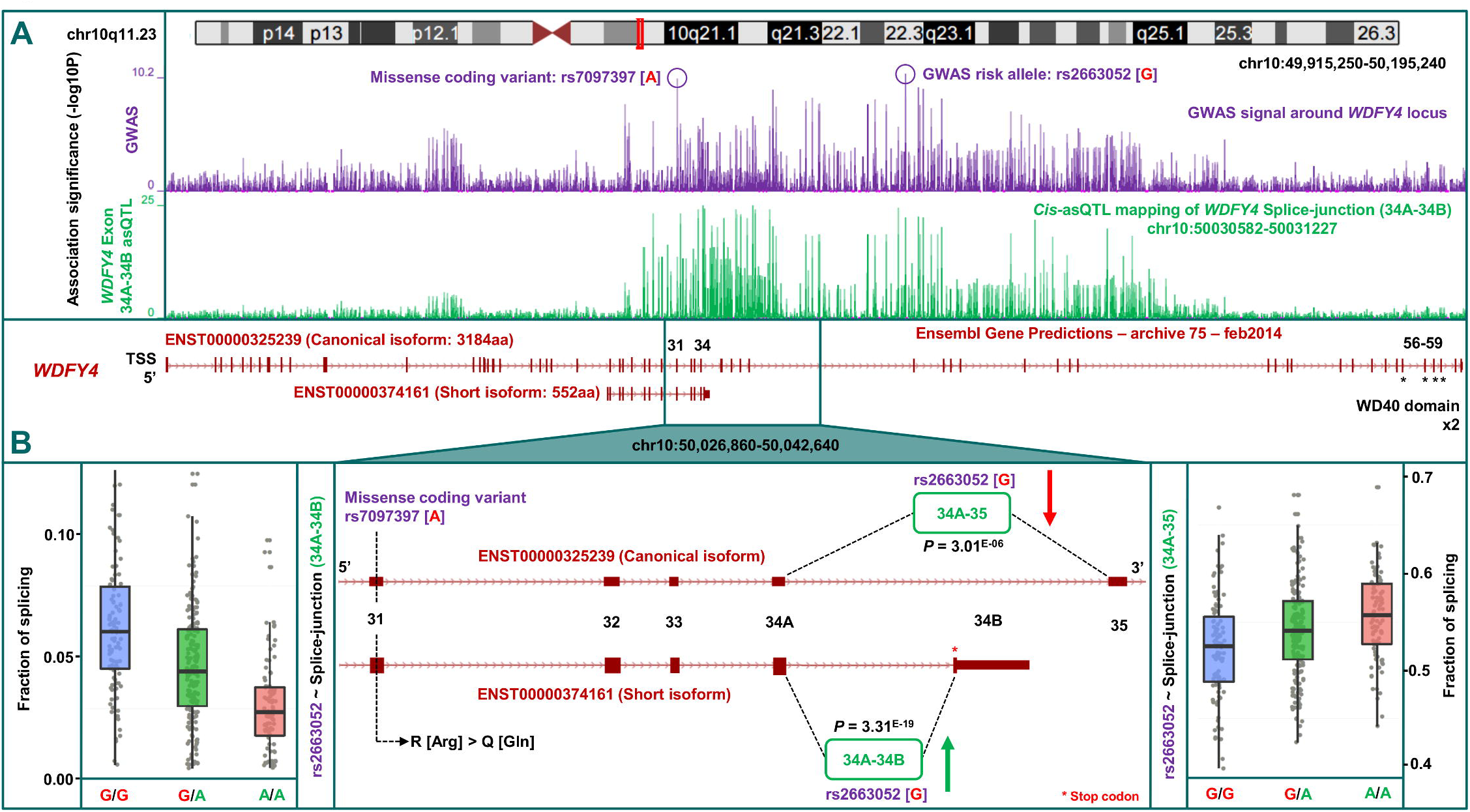
Identification of potential disease-associated mechanisms at the *WDFY4* susceptibility locus using asQTL mapping. **A**) Our SLE GWAS indicates *WDFY4*as the candidate gene at the chr10q11.23 locus tagged by intronic variant rs2663052, as well as the missense coding variant rs7097397 in exon 31 that is in strong LD. *Cis*-eQTL showed rs2663052 is correlated with upregulation of the exon 34A–34B junction of *WDFY4* (signal is colocalised with GWAS) that is unique to the short isoform (ENST00000374161). This isoform lacks the two enzymatic WD40 domains of the full length isoform (ENST00000325239). **B**) Two potential functional mechanisms may occur when harbouring the risk haplotype that carries both risk alleles. Firstly an Arg to Gln amino-acid substitution by rs7097397 in exon 31 that is shared by both the canonical and short isoforms of *WDFY4*, and secondly an upregulation of the short isoform (*P*=3.31^E-19^)that lacks functional domains, caused by rs2663052 or correlated variants, with corresponding down-regulation of the full-length isoform (*P*=3.01^E-06^).

#### IRF5

The GWAS SNP and known asQTL, rs3757387, causes differential promoter usage of *IRF5* (Interferon regulatory factor 5); a molecular mechanism that has previously been reported in predisposition of SLE (80). Alteration of a consensus splice-site causes upregulation of a shorter isoform of *IRF5* which subsequently leads to erroneous activation of the type-1 IFN-pathway and pro-inflammatory cytokines (81). We replicated this known effect by observing an increased fraction of splicing of the shorter isoform of *IRF5*, ENST00000489702, with respect to the risk allele rs3757387[C] (**Table 8**). The risk allele increases splicing from the first exon to the penultimate exon of ENST00000489702 (*P*=2.2×10^-08^); and from the first exon to the final exon of ENST00000489702 (*P*=5.9 ×10^-07^).

#### NADSYN1

Finally, using splice-junction quantification, we were able to pinpoint the specific transcript of *NADSYN1* that drives the exon-level association previously described (**Table 5**, **Fig. 3**). The GWAS SNP rs3794060 leads to substantial upregulation of the meta-exon 10 to meta-exon 12 splice-site (*P*=8.0×10^-12^) which is unique the ENST00000528509 transcript of *NADSYN1* (**Table 8**, **S14 Fig.**). As a consequence of this splicing event, it appears the meta-exon 11 to meta-exon 12 splice-site is highly reduced (*P*=2.1×10^-14^) with reference to the risk allele [C]. Meta-exons 11 and 12 were implicated in exon-level analysis but a specific transcript could not be isolated as the gene annotation was not collapsed to the granularity used in the asQTL analysis. Interestingly, transcript ENST00000528509 is translated to a 294 amino acid residue protein where the canonical transcript of *NADSYN1*, ENST00000319023, is 706 amino acids. The shorter protein lacks the NAD(+) Synthetase domain (located in positions 339-602aa) found in the canonical protein (Pfam: PF02540); thus implicating loss of this domain as a potential causal mechanism.

## Discussion

Detailed characterization of the functional effects of human regulatory genetic variation associated with complex-disease is paramount to our understanding of molecular aetiology and poised to make significant contributions to translational medicine (82). Use of eQTL mapping studies to interpret GWAS findings have proved fundamental in our progression towards this goal - through prioritization of candidate genes, refinement of causal variants, and illumination of mechanistic relationships between disease-associated genetic variants and gene expression (82, 83). However, there is often a disparity between disease-associated genetic variation and phenotypic alteration, which historically may be due to the use of microarray-based technologies to profile genome-wide gene expression. With the advent of RNA-Seq, we can achieve more accurate quantification of the mRNA output of genes, individual exons, and isoform abundance, as well as unannotated and non-coding transcripts. Detection of splicing variants at susceptibility loci using RNA-Seq has the potential to uncover the role of specific isoforms implicated in disease risk, which are likely to have remained concealed by microarray, as a largely independent subset of variants control alternative splicing of isoforms compared to overall gene abundance (35).

Our two major motivations for this study were firstly to directly compare the ability of RNA-Seq with microarrays to detect candidate-causal eQTLs and their associated eGenes from GWAS data, and then to assess each platform’s effectiveness in explaining the potential causal genes and mechanisms implicated by SLE risk alleles. A previous investigation of the same SLE risk alleles used in this study with eQTL data from LCL microarray datasets revealed that only 13 of the 39 risk alleles were eQTLs for a total of 15 eGenes (**Table S6**). Therefore, in an attempt to increase our understanding of the extent to which eQTLs can explain the functional consequences of our risk alleles and to achieve both of our major aims for this investigation, we set up an analytical pipeline to compare the results of eQTL annotation for 39 SLE susceptibility loci with *cis*-eQTL data from both microarray and RNA-Seq experiments from the TwinsUK cohort in LCLs. We incorporated steps to minimize false-positive associations through conditional and colocalisation analysis, as approximately 34% of all genes have a second independently associated *cis*-eQTL for any of their exons when conditioning on the best *cis*-eQTL (35). The analytical pipeline we present will be applicable to the functional annotation of susceptibility loci from a wide range of human diseases.

**Fig. 7** summarises the data generated in this manuscript illustrating which of the GWAS SNPs show evidence of a candidate-causal eQTL association across the quantification types. Our data analysis revealed that RNA-Seq is a more powerful in the identification of candidate-causal eQTLs and their accompanying eGenes than corresponding microarray datasets (**S6 Table**). Many of the published SLE candidate eGenes and associated mechanisms were well-replicated by performing *cis*-eQTL analysis of RNA-Seq datasets at various resolutions for each of the GWAS variants. These eGenes included, among others, the effect of risk variants at *BLK* and *FAM167A* (**S9 Fig.**), *MIR146A* (**Fig. 4A**), *BANK1* (**S12 Fig.**) and *IRF5*. Microarray studies were unable to detect the novel SLE candidate-causal eGenes identified from RNA-Seq data including *NADSYN1, TCF7*, *SKP1*, *WDFY4*, *IKZF2*, and the non-coding RNA genes: *RP11-148O21.2, RP11-148O21.4*, and *RP11-660L16.2* (**Fig. 2**).

**Fig. 7:**
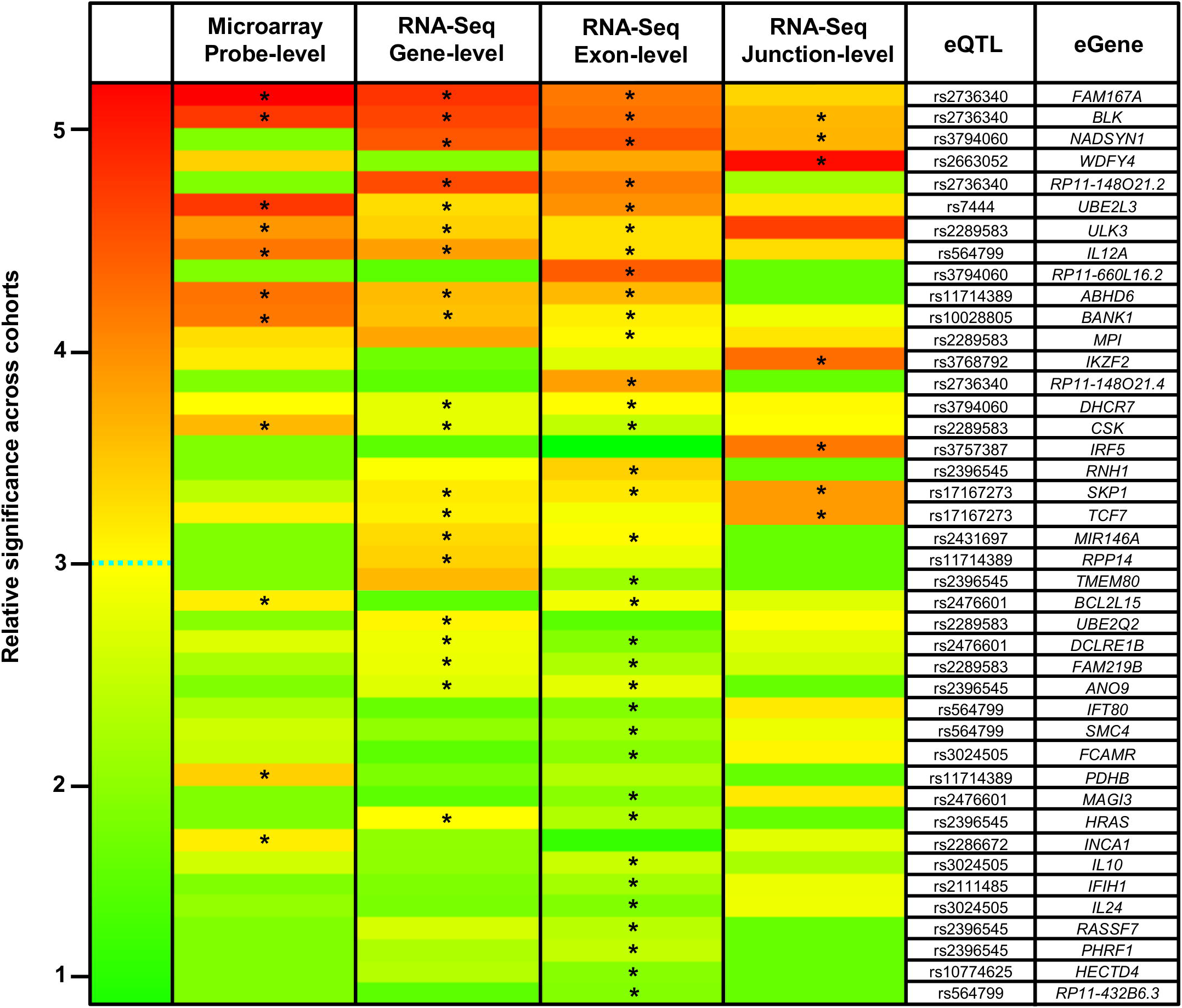
Heatmap of candidate-causal eQTLs and eGenes detected across all expression-quantification types. Heatmap of all candidate-causal *cis*-eQTL associations across the four quantification types (microarray, RNA-Seq gene-level, RNA-Seq exon-level, and RNA-Seq splice-junction level). The first column is the key showing the relative *P* value of the eQTLs within each platform. For the platform-specific columns, if an eQTL-eGene association is candidate-causal in at least one quantification type, the data is displayed across all platforms. Rows are or ordered by decreasing cumulative significance across quantification types. To normalize across quantification types, relative significance of each association per column was calculated as the –log_2_ (*P*/*P*_max_); where *P*_max_ is the most significant association per quantification type. If an association is deemed to be candidate-causal within a particular profiling-type, it is highlighted with an asterisk.

Our results also demonstrate that RNA-Seq analysis is much better than microarrays in identifying multiple eGenes for a single SNP (that may tag multiple functional variants). An increased ratio of eQTLs to eGenes (average number of eGenes per candidate-causal eQTL) was observed using RNA-Seq at exon-level (2.42) compared with gene-level (1.72) and which were both greater than microarray (1.5) (**S5 Fig.**). The ability of RNA-Seq exon-level analysis to identify multiple target eGenes for a specific eQTL is supported by recent observations from capture Hi-C (cHi-C) experiments to functionally annotate chromatin interactions, such as enhancer–promoter interfaces (84, 85). It has been shown that chromatin interactions can control transcription in both *cis* and *trans* in a largely sequence-specific manner, thus it is likely that some GWAS variants may functionally act through the disruption of chromatin dynamics resulting in perturbation of expression of multiple genes (84, 86, 87). Specific instances of this type of effect are seen in colorectal cancer risk loci where the risk locus 11q23 mapped to interactions with genes *C11orf53*, *C11orf92* and *C11orf93,* and separately, the risk SNP rs6983267 within 8q24 disrupts a chromatin regulatory network involving interactions between three genes *CCAT2*, *CCAT1* and *MYC* (84). Our results support this notion of multiple perturbed genes at a single susceptibility locus. At 1q32.1, for example, rs3024505 was found to be associated with three plausible candidate-causal eGenes: *IL10*, *IL24*, and *FCAMR* (located 1 kb, 130 kb, and 191 kb away from rs3024505 respectively). These chromatin capture data also support the argument of using RNA-Seq and extending the *cis*-eQTL distance (typically +/-0.25–1Mb) to a larger region (+/-5Mb) around the associated SNP to identify effects caused by chromatin interactions over a larger distance than commonly designated as *trans* from eQTL-type analyses (88). Integration of eQTL data with epigenetic regulation (promoter methylation, histone modification and expression of non-coding RNA) will allow the identification of the potential mechanism of action (disruption of epigenetic landscape) and disease biology of associated variants.

We have also demonstrated the power of RNA-Seq compared with microarray in the discovery of alternative-splicing events. This is of significant importance as approximately 80% of all human genes undergo alternative splicing and it is estimated that 20-30% of disease-associated mutations modify the configuration of expressed isoforms (89, 90). A recent study has concluded that regulatory variants controlling gene splicing are major contributors to complex traits (91). Two examples of this taken from our analyses are the risk alleles at the *IKZF2* and *WDFY4* loci which drive up-regulation of short isoforms. Neither of these effects were captured by the microarray probes that targeted the 3′-UTR of the canonical longer isoforms. At *WDFY4*, the splicing variant rs2263052[G] is in strong LD with an (Arg1816Gln) missense variant rs7097397[G], which we showed to be associated with SLE (**Fig. 6**). These two potentially causal signals may reinforce each other. The novel association identified in our group’s recent GWAS study (63) at the *IKZF2* locus implicated a risk haplotype tagged by the risk allele of rs3768792[G]. We identified, by RNA-Seq splice-junction quantification exclusively, that variation at the rs3768792[G] risk allele led to increased production of a shorter isoform of *IKZF2* (**Fig. 5**). Interestingly, other members of this gene family, *IKZF1* and *IKZF3*, are also associated with SLE (63). An associated variant in the 3′-UTR of *IKZF1* has been associated with increased expression of genes of the type 1 IFN pathway and decreased expression of complement genes, both mediated in *trans* (64). The functional effect of the *IKZF3* common variant is less well documented, although *IKZF3* knockout mice develop spontaneous autoantibodies and B-cell lymphoma (92). The *IKZF2* knockout mouse has not been characterised for immune-deficient phenotypes. We hypothesize that upregulation of the shorter isoform of *IKZF2* caused by rs3768792[G], which lacks the dimerization domain, reduces translocation of the protein into the nucleus and regulation of transcription of target genes. We believe this aberrant mechanism may result in loss of T-reg stability. The asQTL discoveries described in this manuscript are examples of how RNA-Seq can suggest a potential causal mechanism that can be easily validated experimentally.

Our study also demonstrates that RNA-Seq is able to identify disease-relevant non-coding RNAs. These type of transcripts have long been known to be of relevance to human disease, however their detection and functional importance may have been under-estimated. We identified three novel non-coding anti-sense RNAs using RNA-Seq: two regulated by rs2736340 (*RP11-148O21.2* and *RP11-148O21.4*) (**Fig. 4B**), and one regulated by rs3794060 (*RP11-660L16.2*) (**Fig. 4C**). These findings were replicated in whole blood (**Table 7**).

The data we present in this manuscript demonstrate that a comprehensive integrated approach for eQTL analysis should be undertaken at gene-, exon- and exon-junction level quantification. Excluding one or more levels of analysis will mean that eQTLs and/or eGenes may be missed. This is illustrated by a number of examples in **Fig. 7**. At some loci, there was an exon-level effect, which was not observed at the gene-level. For example, the risk allele rs3024505[A] was an eQTL for *IL10*, *IL24*, and *FCAMR* at exon-level resolution only (**Table 5**), but none of these eGenes were deemed to be significant at gene-level. This suggests that the exon-level effect is more targeted than gene-level quantification, where multiple different effects across the gene may dilute out the signal. However, some eGenes exhibit a probable whole gene-level effect (*UBE2L3*, *BLK*, and *FAM167A)* as every expressed exon showed a candidate-causal association for the respective risk alleles. Hierarchical clustering tests could be designed to distinguish between these genuine gene-level and exon-level effects, such as at *NADSYN1* or *BANK1*, where the gene-level effect is likely to be driven by only a subset of exons (**Fig. 3**, **S12 Fig.**). There were occurrences where significant candidate-causal eGenes were detected but there was no effect at the exon-level. At *TCF7*, variants in low LD with rs7726414 exhibited significant exon-level *cis*- eQTLs (**S3 Table**) and were deemed to be independent of the disease-association. However, gene-level analysis revealed that the risk rs7726414 variant was candidate-causal for total *TCF7* expression (**S2 Table**). These results emphasise that for any given eGene there may be multiple genetic effects at different resolutions of quantification.

We understand the limitation of LCLs for transcript profiling studies. There is an inherent limitation analysing LCL expression datasets, such as those available for our study, because although LCLs are a good surrogate model for primary B-cells, the effect of EBV transformation is likely to disrupt their underlying epigenetic and transcriptomic background. The percentage of asQTLs in LCLs will exhibit significantly less replication in primary cell types because of cell-type variability in the genetic control of isoform usage, approximately 70% of *cis*-eQTL detected in LCLs can be replicated in a primary tissue type (33). The significance of alternative-splicing in genomic medicine will become better understood once large RNA-Seq based eQTL cohorts emerge across a multitude of disease-relevant cell-types. A gold standard of candidate-causal eQTL mapping strategies using RNA-Seq across datasets using an explicit set of quantification types (gene-, exon-, splice-junction, isoform) and analytical pipelines, will accelerate this process.

In summary, we have demonstrated the effectiveness of eQTL analysis using RNA-Seq by increasing the numbers of candidate eGenes regulated by SLE associated alleles (**Fig. 2**, **Fig. 7**, and **S6 Table**). We have shown that the power of RNA-Seq in eQTL annotation of GWAS loci lies not only the assessment of the variants regulating the expression of candidate genes, but also in uncovering putative molecular mechanisms, which allow for more refined targeted follow-up studies to assess the phenotypic consequence of the disease-associated variant. These studies could include knocking down target gene or introducing recombinant vectors *in vitro* to overexpress target genes to evaluate phenotypic consequence of expression changes of specific loci. Site directed mutagenesis could be used to introduce candidate causal splice-sites and over-express target isoforms. The CRISPR/Cas9 system for targeted genome editing presents an exciting opportunity for eQTL/RNA-Seq targeted follow-up studies and the investigation of the effect that specific variants have on the expression profile across different cell types.

## Materials and Methods

### Selection of SLE associated SNPs

SLE associated SNPs were taken from our recent 2015 publication (63). The study comprised a primary GWAS, with validation through meta-analysis and replication study in an external cohort (7,219 cases, 15,991 controls in total). The independently-associated susceptibility loci taken forward for this investigation were those that passed either genome-wide significance (*P*< 5×10^-08^) in the primary GWAS or meta-analysis and/or those that reached significance in the replication study (False Discovery Rate, q<0.01). We defined the ‘GWAS SNP’ at each locus as either being the SNP with the lowest *P*-value post meta-analysis or the SNP with the greatest evidence of a missense effect as defined by a Bayes Factor. We omitted non-autosomal associations and those within the Major Histocompatibility Complex (MHC), and SNPs with a MAF <0.05. In total, 39 GWAS SNPs were taken forward (**Table 1**).

### TwinsUK cohort eQTL datasets

Expression profiling by microarray (9) and RNA-Seq (39) of individuals from the UK Adult Twin Registry (TwinsUK) was carried out in two separate studies on the MuTHER (Multiple Tissue Human Expression Resource) cohort (**Table 2**). The MuTHER cohort is composed of 856 healthy female individuals of European descent aged between 37-85 years. We considered expression quantification data from both resting LCLs and whole blood. Profiling by microarray was performed using the Illumina Human HT-12 V3 BeadChips. For RNA-Seq, samples were sequenced using the Illumina HiSeq2000 and the 49-bp paired-end reads mapped with BWA v0.5.9 to the GRCh37 reference genome. Exons (‘meta-exons’ created by merging all overlapping exonic portions of a gene into non-redundant units) were quantified using read-counts against the GENCODE v10 annotation; with gene quantification defined as the sum of all exon quantifications belonging to the same gene. Full quality control and normalization procedures are described in the respective articles. Data from each of the TwinsUK eQTL studies (**Table 2**) were provided in different formats. In each instance it was necessary to generate summary *cis*-eQTL statistics per GWAS SNP (SNP, expression-unit, β, standard error of β, and *P*-value of association) for integration analysis. Per quantification type (microarray, RNA-Seq gene-level, and exon-level), each GWAS SNP was subject to *cis*-eQTL analysis against all expression-units within +/-1Mb using no *P*-value threshold. If the GWAS SNP was not found in an eQTL dataset, the most highly correlated, closest tag SNP with r^2^ ≥ 0.7, common to all datasets, was used as its proxy (**Table 1**). Adjustment for multiple testing of *cis*-eQTL results per quantification type were undertaken using FDR with q <0.05 deemed significant.

#### Microarray cis-eQTL mapping

We used the Genevar (GENe Expression VARiation) portal to generate summary-level *cis*- eQTL results (50). We ran the association between normalized expression data of the 777 individuals and each GWAS SNP implementing the external algorithm option (two-step mixed model–based score test). In total 768 probes (559) genes, were tested.

#### RNA-Seq (gene-level) cis-eQTL mapping

RNA-Seq gene-level quantification was provided as residualized read-counts (effect of family structure and other covariates regressed out). We had full genetic data for 683 individuals and performed the analysis of each GWAS SNP against the transformed residuals using the linear-model function within the MatrixeQTL R package (93). 520 genes were tested against in *cis*.

#### RNA-Seq (exon-level) cis-eQTL mapping

*P*-values from the association of all SNPs against exon-level quantifications for 765 individuals using linear-regression were provided. We generated the t-statistic using the lower-tail quantile function t-distribution function in R with 763 degrees of freedom. The standard error and β were derived from the t-statistic. We then extracted the summary *cis*- eQTL results for each GWAS SNP. 4,786 exons, corresponding to 716 genes for testing.

### SLE candidate-causal *cis*-eQTL classification

#### Conditional analysis

We used the COJO (conditional and joint genome-wide association analysis) function of the GCTA (Genome-wide Complex Trait Analysis) application to determine whether the GWAS SNP had an independent effect on expression from that of the best *cis*-eQTL (55). For each significant association (q<0.05), we re-performed the analysis using all SNPs within +/-1Mb of the expression-unit in hand. We used the available genotype information of the 683 TwinsUK individuals to extract allele coding along with the MAF, and integrated this with the *cis*-eQTL summary data. We discarded SNPs with: MAF < 0.05, imputation call-rates < 0.8, and HWE *P*<1x10^-04^. We used these individuals as the reference panel to calculate local pairwise linkage disequilibrium (LD) between variants. Per significant association, all *cis*- eQTLs were conditioned on by the best *cis*-eQTL. We then extracted the conditional *P*-value of the GWAS SNP and considered associations to be independent to the best *cis*-eQTL if P_cond_<0.05.

#### Colocalisation Analysis

We employed the 'coloc' Bayesian statistical method using summary data implemented in R to test for colocalisation between *cis*-eQTL and disease causal variants derived from the GWAS (56). The method makes the assumption of there being a single causal variant for each trait (disease association and gene-expression from two separate studies) per locus and calculates the posterior probabilities under five different causal variant hypotheses: association with neither trait (H0), association with one trait but not the other (H1, H2), association with both traits but from independent signals, and association with both traits with a shared causal signal (H4). We extracted the necessary SNP statistics for the disease-associated regions from our own GWAS and applied the same SNP filters used in the conditional analysis. We tested for colocalisation between the GWAS summary data and *cis*- eQTL data for each significant association within a +/-1Mb window of the GWAS SNP. We assigned the prior probabilities, p1 and p2 (SNP is associated with GWAS and gene expression respectively), as 1 × 10^-04^ i.e. 1 in 10,000 SNPs are causal to either trait, with p12 (SNP is associated with both traits) as 1×10^-06^ or 1 in 100 SNPs associated with one trait are also associated with the other. For each *cis*-eQTL association colocalisation test, if the posterior probability PP3 (two distinct causal variants, one for each trait) is greater than PP4 (single causal variant common to both traits), then greater posterior support is given to the hypothesis that independent causal variants exist in both traits and thus the eQTL is unlikely to be attributed to SLE genetic association.

### Definition of SLE candidate-causal cis-eQTL and eGene

We defined a GWAS SNP as an SLE candidate-causal *cis*-eQTL if it met the following criteria: significant post-multiple testing adjustment (q < 0.05), not independent to the best *cis*-eQTL from conditional analysis (*P*_cond_ > 0.05), and supporting evidence of a shared causal variant between gene expression and the primary GWAS signal based on colocalisation (PP3 < PP4). The gene whose expression is modulated by the candidate-causal eQTL is defined as an SLE candidate-causal eGene (**Fig. 1**).

### Validation of LCL SLE candidate-causal cis-eQTLs in whole blood

*Cis*-eQTL summary data from whole blood at RNA-Seq exon-level were made available for 384 individuals of the 856 TwinsUK cohort individuals (**Table 2**). Expression profiling and genotyping were identical to that as described for LCLs. We applied the same methodology to this dataset to generate full *cis*-eQTL summary statistics, perform conditional and colocalisation analysis, and classify SLE candidate-causal eQTLs and associated eGenes (**Fig. 1**). In total, 3,793 exons were tested against, corresponding to 654 genes.

### Geuvadis SLE candidate-causal *cis*-asQTL analysis

We investigated SLE disease-associated alternative splicing QTLs (asQTLs) using European samples from the raw alignment files of the Geuvadis (35) 1000 Genomes RNA-Seq project profiled in LCLs (**Table 2**). Genotype data and read-alignments were downloaded from ArrayExpress for the 373 Europeans (comprising 91 CEU, 95 FIN, 94 GBR, and 93 TSI). We performed PCA on chromosome 20 using the R/Bioconductor package SNPRelate (94) and decided to include the first three principle components as covariates in the eQTL model as well as the binary imputation status (mixture of Phase 1 and Phase 2 imputed individuals). We removed SNPs with MAF<0.05, imputation call-rates<0.8, and HWE *P*<1×10^-04^. We removed non-uniquely mapped, non-properly paired reads, and reads with more than eight mismatches for read and mate using Samtools(95). We used the Altrans (96) method against GENCODE v10 to generate relative quantifications (link-counts) of splicing events; which in brief, utilizes split and paired-end reads to count links between exon-boundaries, which themselves are created by flattening the annotation into unique non-redundant exon-groups. Following PCA of the link-counts, we decided to normalize all link-counts with the first 10 principle components then removed exon-boundaries with zero links in more than 10% of individuals. Link-counts were converted to link-fractions (coverage of the link over the sum of the coverage of all the links that the first exon makes) and merged in both 5′-3′ and 3′-5′ directions. Per GWAS SNP we performed *cis*-eQTL analysis against the normalized link-fractions in MatrixeQTL with a linear-model (93). 33,039 link-fractions were tested against corresponding to 817 genes in total. After FDR multiple-testing adjustment we considered associations with q<0.05 as significant. As full genetic and expression data were available, we decided to use the Regulatory Trait Concordance (RTC) method to assess the likelihood of a shared functional variant between the GWAS SNP and the asQTL signal (48). For each significant asQTL association we extracted the residuals of the linear-regression of the best *cis*-eQTL against normalized link-fractions and re-performed the analysis using all SNPs within the defined hotspot interval against this pseudo-phenotype. The RTC score was defined as (N_SNPs_-Rank_GWAS SNP_)/N_SNPs_ where N_SNPs_ is the number of SNPs in the interval, and Rank_GWAS SNP_ is the rank of the GWAS SNP association *P*-value against all other SNPs in the interval. We classified an SLE candidate-causal *cis*-asQTL as a GWAS SNP with a significant association (q < 0.05) with link-fraction quantification and an RTC score > 0.9.

### Statistical analysis and data visualisation

We performed statistical analysis, graphics and data handling in R version 3.2.0 and ggplot2. Genetic plots were generated using LocusZoom v1.1 (97). Karyotype diagrams were modified from Ensembl (98). GWAS association plots and gene annotation graphic visualisations were created using the UCSC Genome Browser(99).

## Acknowledgements

DSCG and TJV were awarded an Arthritis Research UK PhD Studentship (for CO) (20332) “SLE pathogenesis: the molecular basis for multiple association signals in *IKZF2*” to undertake the analyses described in this manuscript. DSCG, DLM and TJV were awarded an Arthritis Research UK Project Grant (20265) “Post GWAS: A Functional Genomic Analysis of the SLE Susceptibility Gene *IKZF1*”, which supported AC. TJV received funds from China Scholarship Council, number 201406380127 for LC. ALR was supported by MRC project grant L002604/1 “Functional Genomics of SLE: A Trans-ancestral approach” and an ARUK project grant (20580) “Targeted DNA sequencing in Indian and European samples contributes to causal allele identification for lupus”. KSS was awarded funding by the MRC (MR/L01999X/1 and MR/M004422/1).

The Geuvadis 1000 Genomes RNA-Seq data was downloaded from the EBI ArrayExpress Portal (accessions E-GEUV-1). The RNA-Seq data on the TwinsUK samples in both LCLs and whole blood was made available through the EUROBATS project (EGAS00001000805 from the European Genome-Phenome Archive). The expression microarray data on the TwinsUK samples was downloaded from the EBI ArrayExpress Portal (E-TABM_1140).

The TwinsUK study was funded by the Wellcome Trust; European Community’s Seventh Framework Programme (FP7/2007-2013). The study also receives support from the National Institute for Health Research (NIHR)- funded BioResource, Clinical Research Facility and Biomedical Research Centre based at Guy's and St Thomas' NHS Foundation Trust in partnership with King's College London. SNP Genotyping was performed by The Wellcome Trust Sanger Institute and National Eye Institute via NIH/CIDR.

## Supporting Information

**S1 Table. All significant eQTLs (q < 0.05) and associated eGenes detected at microarray (probe-level) with conditional and colocalisation results.**

**S2 Table. All significant eQTLs (q < 0.05) and associated eGenes detected at RNA-Seq (gene-level) with conditional and colocalisation results.**

**S3 Table. All significant eQTLs (q < 0.05) and associated eGenes detected at RNA-Seq (exon-level) with conditional and colocalisation results.**

**S1 Fig. Number of eQTL discoveries per quantification type.** Including significant associations (q<0.05), and candidate-causal associations (significant, and not-independent and colocalised with GWAS).

**S2 Fig. Number of eGene discoveries per quantification type.** Including significant associations (q<0.05), and candidate-causal associations (significant, and not-independent and colocalised with GWAS).

**S3 Fig. Shared candidate-causal eQTLs per quantification type.**

**S4 Fig. Shared candidate-causal eGenes per quantification type.**

**S5 Fig. Ratio of eQTLs to candidate-causal eGenes per quantification type.**

**S6 Fig. Gene-level and exon-level candidate-causal associations with *TCF7* and *SKP1* against rs7726414.** *Cis*-eQTL analysis at gene-level and exon-level using RNA-Seq implicate novel SLE-associated eGenes *TCF7* and *SKP1*.

**S7 Fig. High gene-density over associated variants tagged by GWAS SNP rs12802200.** The six candidate-causal eGenes against rs1280220 discovered using RNA-Seq at either quantification method are marked with an asterisk.

**S8 Fig. Candidate-causal eGenes *DHCR7* and *NADSYN1* for rs3794060, and non-coding eGene *RP11-660L16.2***. The GWAS SNP rs3794060 is a candidate-causal eGene for *DHCR7* and *NADSYN1*, and also the non-coding eGene *RP11-660L16.2* at exon-level; which is located between *DCHR7* and *NADSYN1*.

**S9 Fig. Candidate-causal eGenes *FAM167A*, *BLK* and two non-coding RNAs (*RP11- 138021.4* and *RP11-138021.2*) driven by rs2736340.** Associated variant rs2736340 lies in a region of intense regulatory chromatin marks located in the bi-directional promoter of *BLK* and *FAM167A* which are both candidate-causal eGenes at RNA-Seq gene-level. At exon-level, the non-coding eGenes *RP11-138021.4* and *RP11-138021.2* are also candidate-causal.

**S10 Fig. Effect-size correlation of GWAS SNP associations with matched *cis*-exons between LCL and whole-blood.**

**S4 Table. All significant eQTLs (q < 0.05) and associated eGenes detected at RNA-Seq (exon-level) l with conditional and colocalisation results in whole-blood.**

**S11 Fig. Candidate-causal eQTLs and eGenes in whole blood.** Comparison with LCL associations.

**S12 Fig. Whole-blood exon-level eQTL effect on *BANK1* exon 2.** Correlation between GWAS SNP and the best whole-blood eQTL for *BANK1* exon 2. Both are highly correlated with known branch-point SNP. All are weakly correlated with best LCL eQTL for *BANK1* exon 2.

**S13 Fig. Exon-level eQTL analysis of *NADSYN1* in whole-blood and LCLs reveal near-identical splicing effect.** Meta-exons 11 and 12 are substantially disrupted with reference to GWAS SNP rs3794060 in both LCLs and whole-blood.

**S5 Table. All significant asQTLs (q < 0.05) and associated eGenes detected at RNA-Seq (splice-junction level).**

**S14 Fig. Proposed splicing mechanism of *NADSYN1* caused by risk haplotype tagged by rs3794060.** *NADSYN1* Ensembl transcript annotation displayed. *Cis*-asQTL identified the meta-exon 10 to meta-exon 12 junction is upregulated with risk allele [C] and consequently the meta-exon 11 to meta-exon 12 junction is downregulated.

**S6 Table. Comparison of eQTLs and eGenes for SLE risk alleles between previously reported in microarray studies and from RNA-Seq in current study.**

## Abbreviations

GWAS: Genome-Wide Association Study
eQTL: expression Quantitative Trait Loci
RNA-Seq: RNA-Sequencing
SLE: Systemic Lupus Erythematosus
asQTL: alternative-splicing Quantitative Trait Loci
SNP: Single Nucleotide Polymorphism
LCL: Lymphoblastoid cell line

